# Restoration of Keratinocyte Homeostasis Drives Resolution of Skin Inflammation

**DOI:** 10.64898/2026.03.04.708224

**Authors:** Dennis Gruszka, Rachael Bogle, Roopesh Singh Gangwar, Brian Richardson, Josh Webster, Jennifer Fox, Michele Mumaw, Athira Sivadas, Mark J. Cameron, Thomas S. McCormick, Lam C. Tsoi, Johann E. Gudjonsson, Nicole L. Ward

**Affiliations:** Department of Dermatology, Case Western Reserve University, Cleveland, Ohio; Department of Nutrition, Case Western Reserve University, Cleveland, Ohio; Department of Dermatology, Vanderbilt University Medical Center, Nashville, TN; Department of Dermatology, University of Michigan, Ann Arbor, MI; School of Medicine, Vanderbilt University, Nashville, TN; Vanderbilt Institute for Infection, Immunology, and Inflammation, Vanderbilt University Medical Center, Nashville, TN; Vanderbilt Center for Immunobiology, Vanderbilt University Medical Center, Nashville, TN

## Abstract

Chronic skin inflammation is sustained by reciprocal interactions between epidermal dysfunction and immune activation, yet whether epithelial state actively governs restoration of tissue homeostasis remains unclear. Using a murine model of inflammatory skin disease, we modulated epidermal lipid metabolism and examined its effects on tissue organization. Transcriptomic profiling revealed coordinated reversal of inflammatory, metabolic, and structural gene programs accompanied by normalization of epidermal architecture. Single-cell RNA sequencing showed that this remodeling was concentrated in differentiated keratinocytes, with suppression of IL-17 and neutrophil-associated responses and restoration of barrier and mitochondrial-lipid programs, while stromal and myeloid compartments displayed secondary adaptation. Cross-species analysis demonstrated that resolution-associated gene networks are inversely regulated in human psoriasis. Integrated proteomic and transcriptomic analyses further identified a conserved epithelial regulatory triad whose concordant regulation in psoriasis and atopic dermatitis, and whose in vivo silencing, establish mechanistic control of disease severity. Together, these findings indicate that inflammatory resolution reflects reorganization of epidermal transcriptional networks and position epithelial state as a determinant of inflammatory persistence.

## Introduction

Barrier tissues such as skin continuously integrate structural integrity, metabolic demands, and immune surveillance. In this context, epithelial cells are not passive substrates for inflammation but active regulators of tissue homeostasis, capable of sensing environmental cues and shaping immune behavior ^1, 2, 3^. Disruption of epithelial differentiation, lipid metabolism, ion handling, or mitochondrial function can destabilize barrier integrity and amplify inflammatory loops, whereas restoration of epithelial competence may enable intrinsic resolution programs to engage. Although these concepts are increasingly recognized, direct mechanistic evidence that epithelial state alone can impose inflammatory quiescence in established disease remains limited.

Dietary long-chain ω-3 polyunsaturated fatty acids (PUFAs) represent a physiologically relevant system-level intervention, identified a priori by a mechanism-based prediction framework (mmPredict), and used here to interrogate epithelial regulatory capacity in inflamed skin ^4, 5^. Rather than acting as narrowly anti-inflammatory agents, ω-3 PUFAs engage membrane- and lipid-dependent regulatory processes that drive system-level remodeling of cellular metabolic networks and downstream signaling programs across tissues ^6^.

Omics-based studies across metabolic and inflammatory diseases support this systems-level mode of action; however, how these effects are executed within inflamed skin, and whether they preferentially engage epithelial versus immune compartments, remains unclear. In chronic inflammatory skin diseases, including psoriasis and atopic dermatitis, clinical trials of ω-3 PUFA supplementation have produced heterogeneous outcomes across formulations and study designs, underscoring the need for mechanistic frameworks that define where and how resolution-associated programs operate in diseased tissue ^7, 8, 9, 10, 11, 12^.

Addressing these questions requires combining disease-relevant experimental strategies with cell-resolved and causal analyses. Bulk transcriptomic studies capture integrated tissue responses but cannot assign execution to specific cellular compartments, whereas single-cell profiling localizes transcriptional programs without establishing functional necessity. Integrating these approaches with targeted functional manipulation is therefore essential to determine whether epithelial reprogramming merely accompanies inflammatory resolution or actively drives it.

Here, we use dietary ω-3 PUFA exposure as a system-level biological input in the KC-Tie2 mouse model of chronic skin inflammation ^13, 14, 15, 16, 17^. By integrating bulk and single-cell RNA sequencing, pathway-level analyses, proteomic data, and in vivo functional gene targeting, we define how inflammatory skin pathology is reorganized across compartments and identify epithelial regulatory nodes that causally control disease severity. This approach enables direct testing of whether restoration of keratinocyte-intrinsic homeostasis is sufficient to suppress pathogenic immune circuits and reveals how epithelial regulatory state governs inflammatory outcome.

## Results

### Metabolic intervention restores epidermal architecture and reduces inflammation

To determine whether altering systemic metabolic state could reverse chronic skin inflammation, KC-Tie2 transgenic mice were maintained on standard chow until 6 weeks of age, when robust cutaneous disease is present, and then transitioned to either control chow or chow supplemented with long-chain ω-3 polyunsaturated fatty acids (PUFA) for an additional six weeks (Fig. 1a). The dietary intervention was well tolerated, with no evidence of weight loss or overt adverse effects.

**Figure 1.**
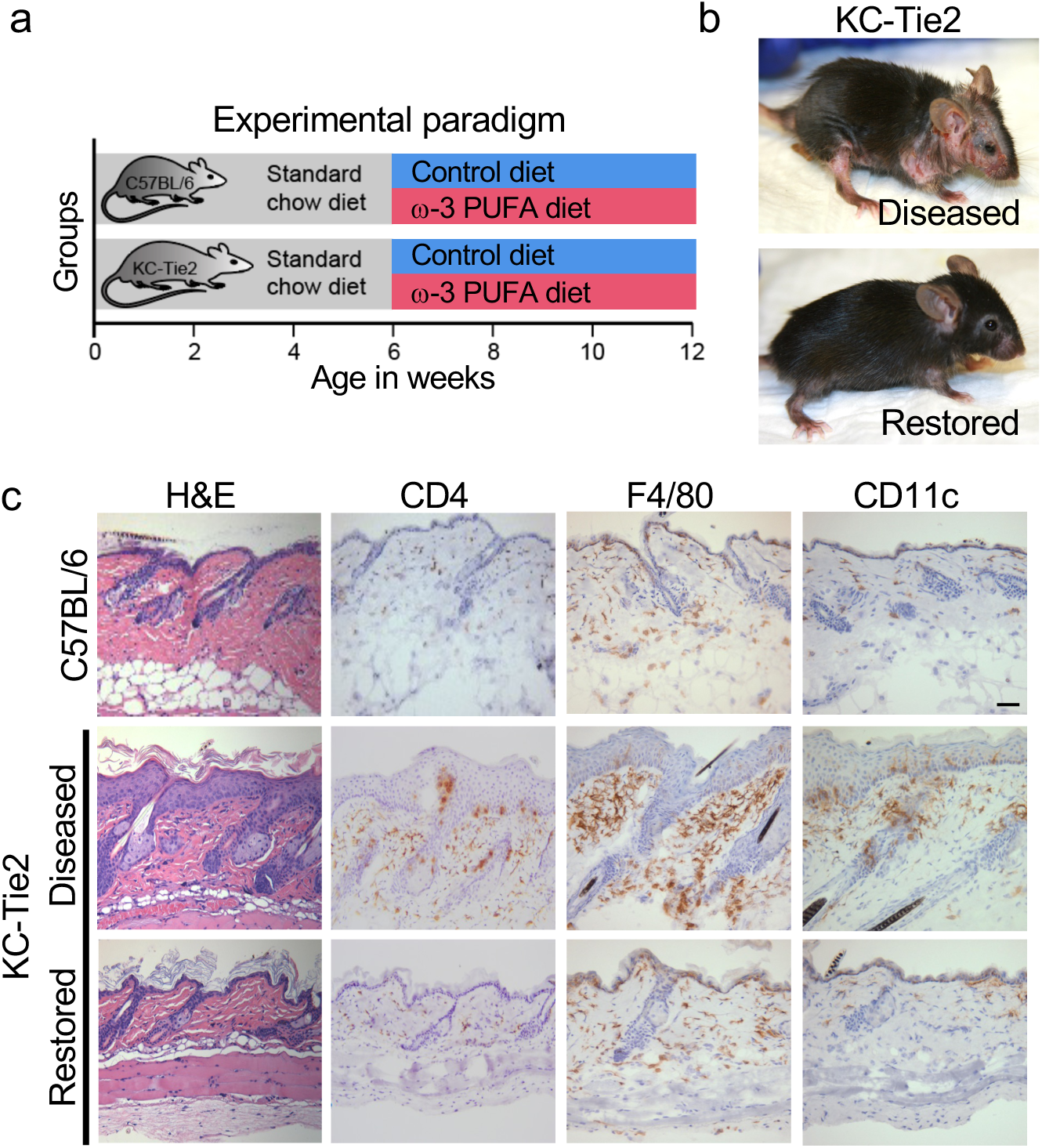
Metabolic intervention restores epidermal architecture in chronically inflamed skin. **a,** Experimental paradigm. C57BL/6 littermates and KC-Tie2 transgenic mice were maintained on standard chow until 6 weeks of age, when psoriasiform skin inflammation is established, and then transitioned to either a control diet or a long-chain ω-3 polyunsaturated fatty acid (PUFA)-enriched diet for an additional 6 weeks. **b,** Representative gross images of KC-Tie2 mice with established disease (*Diseased*) and following intervention (*Restored*). “Restored” denotes KC-Tie2 mice in which gross erythema and scaling are markedly reduced relative to the diseased state and approximate the appearance of wild-type skin. **c,** Histological and immunohistochemical analysis of dorsal skin from C57BL/6 mice, KC-Tie2 mice with established disease (*Diseased*), and KC-Tie2 mice following intervention (*Restored*). Hematoxylin and eosin (H&E) staining demonstrates pronounced epidermal hyperplasia and dermal cellular expansion in diseased KC-Tie2 skin, which are substantially reduced in restored KC-Tie2 skin. Immunostaining for CD4 (T cells), F4/80 (macrophages), and CD11c (dendritic cells) reveals broad attenuation of inflammatory cell accumulation coincident with normalization of epidermal architecture. Scale bar, 50µm. Quantification of epidermal thickness is provided in Supplementary Fig. 1.

Compared with diseased KC-Tie2 littermates maintained on control chow, ω-3 PUFA-exposed KC-Tie2 mice showed clear improvement in gross skin appearance, with reduced erythema and scaling (restored; Fig. 1b). Histological analysis revealed concordant normalization of epidermal and dermal architecture, including marked reduction in epidermal hyperplasia, restoration of stratum corneum organization, and diminished dermal inflammatory infiltrates (Fig. 1c). Quantitative morphometric confirmed a greater than 70% reduction in epidermal thick ness (acanthosis) relative to diseased KC-Tie2 skin (P < 0.0001; Supplementary Fig. 1), approaching levels observed in C57BL/6 controls. In parallel, CD4⁺ T cells, F4/80⁺ macrophages, and CD11c⁺ dendritic cells were significantly reduced compared with diseased controls (Fig. 1c).

These data demonstrate that modifying systemic metabolic state is sufficient to reverse established architectural and inflammatory features of chronic skin inflammation.

### Chronic inflammation reflects a coordinated epidermal transcriptional state

To define the transcriptional programs underlying skin inflammation and its resolution, we performed bulk RNA sequencing on dorsal skin from KC-Tie2 and C57BL/6 littermates under control conditions and following induction of disease resolution. Principal component analysis showed clear genotype-driven segregation, with KC-Tie2 samples diverging significantly from wild-type controls, consistent with a global disease-associated transcriptional state (Supplementary Fig. 2a; Supplementary Tables 1-2). KC-Tie2 samples undergoing disease resolution shifted toward the C57BL/6 transcriptional space, whereas wild-type samples showed minimal separation across conditions, indicating that large-scale transcriptional remodeling is strongly disease dependent. Unsupervised hierarchical clustering recapitulated this pattern, revealing reversal of disease-associated transcriptional dysregulation in KC-Tie2 skin and restoration of expression profiles toward those of healthy controls (Supplementary Fig. 2b). Together, these analyses indicate that established skin inflammation is associated with a coherent, disease-specific transcriptional state that can be broadly normalized at the tissue level.

Differential expression analysis identified 3,713 genes significantly dysregulated in KC-Tie2 skin relative to C57BL/6 controls (|log₂ fold change| ≥ 1.5, FDR < 0.05), defining a robust inflammatory transcriptional signature (Fig. 2a, left; Supplementary Tables 1-3). Genes elevated in disease reflected keratinocyte stress and aberrant differentiation, including cornified envelope and keratinization programs (*Sprr2b, Sprr2d, Krt27, Krt81, Tchh*), together with IL-17-responsive inflammatory mediators and antimicrobial genes (*Il17a, Il17c, Il1f5, Il1f6, Defb3, S100a8/a9*). Pro-inflammatory cytokines including *Il23a, Il17a, Il22, Il1f5, Il1f6*, and *Tnf* were significantly increased in KC-Tie2 skin (Fig. 2b). In contrast, genes associated with epidermal organization, calcium-dependent signaling, and cytoskeletal regulation (*Cacna2d1, Actn3, Myh8, Tnnc1*) were broadly suppressed, consistent with disruption of epithelial structural integrity in inflamed skin.

**Figure 2.**
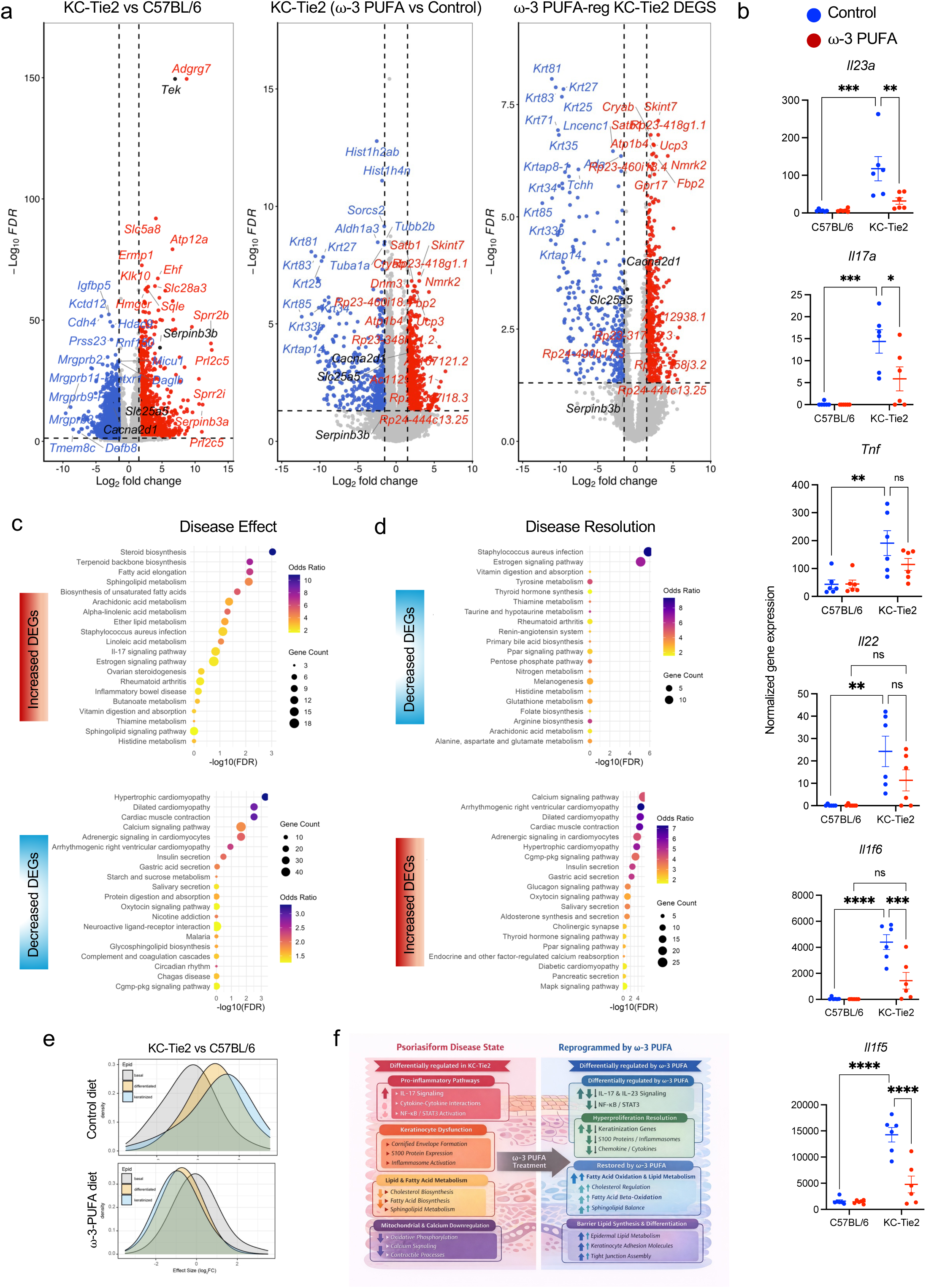
Metabolic intervention reverses disease-associated transcriptional programs. **a,** Volcano plots depicting global differential gene expression across disease and restoration states. **Left:** KC-Tie2 versus C57BL/6 skin under control conditions, defining disease-associated transcriptional changes. **Middle:** KC-Tie2 skin following ω-3 PUFA exposure versus KC-Tie2 control skin, illustrating transcriptional changes accompanying disease normalization. **Right:** Subset of disease-associated genes that are significantly modulated in the opposite direction during ω-3 PUFA-mediated disease resolution. Genes were filtered using |log₂ fold change| ≥ 1.5 and FDR < 0.05; non-significant genes are shown in grey. Selected genes with large effect sizes are labeled. The transgene *Tek* and the epithelial regulators *Serpinb3b*, *Slc25a5*, and *Cacna2d1* are highlighted and examined further in subsequent figures. Summary statistics for all differential expression contrasts, including DEG counts and directionality, are provided in Supplementary Table 3. **b,** Expression of inflammatory mediators (*Il23a*, *Il17a*, *Tnf*, *Il22*, *Il1f6*, and *Il1f5*) in dorsal skin from C57BL/6 and KC-Tie2 mice under control or ω-3 PUFA exposure. Data are shown as individual biological replicates with mean ± s.e.m. Statistical significance was determined by two-way ANOVA with Tukey’s post hoc testing; \**P* < 0.05, \*\**P* < 0.01, \*\*\**P* < 0.001, \*\*\*\**P* < 0.0001; ns, not significant. **c,** KEGG pathway enrichment analysis of genes increased (upper) or decreased (lower) in KC-Tie2 skin relative to C57BL/6 controls (Disease Effect). Enrichment of cardiomyopathy and contractility-related annotations reflect overrepresentation of actin-myosin and calcium-handling genes that are shared across tissues. **d,** KEGG pathway enrichment analysis of genes decreased (upper) or increased (lower) during ω-3 PUFA-mediated restoration in KC-Tie2 skin (Disease Resolution), demonstrating bidirectional normalization of disease-associated pathways. Dot size represents gene count; color denotes odds ratio; the x-axis indicates −log₁₀(FDR). Full gene lists and enrichment statistics are provided in Supplementary Tables 4-7. **e,** Density plots of log₂ fold-change distributions for genes associated with basal, differentiated, and keratinized epidermal compartments. KC-Tie2 versus C57BL/6 skin under control conditions demonstrates disease-associated transcriptional shifts across epidermal layers (upper). ω-3 PUFA-exposed KC-Tie2 skin versus control-fed KC-Tie2 skin shows redistribution of effect sizes toward baseline (lower), indicating that both disease-associated dysregulation and its normalization are concentrated within epidermal gene programs. **f,** Schematic summary integrating transcriptional analyses across disease and resolution states. Chronic inflamed skin is characterized by coordinated activation of IL-17-driven inflammatory signaling, keratinocyte stress responses, disrupted lipid and fatty acid metabolism, and altered calcium and cytoskeletal programs. Disease resolution is associated with suppression of inflammatory modules alongside reactivation of epidermal differentiation, barrier formation, lipid metabolism, calcium signaling, and structural homeostasis, consistent with keratinocyte-centered, systems-level transcriptional reprogramming.

During resolution, 1,354 transcripts were significantly modulated in KC-Tie2 skin (Fig. 2a, middle; Supplementary Tables 1-3). Intersection of disease-associated and resolution-responsive gene sets identified a core group of transcripts whose expression shifted toward control levels (Fig. 2a, right). This group included coordinated suppression of stress and inflammatory genes elevated in disease (e.g., *Sprr2* family members, *Krt81, Il17a, Il17c*), alongside restoration of genes linked to epithelial organization, signaling, and metabolic competence (e.g., *Cacna2d1, Cryab, Gpr17*). Consistent with this shift, *Il23a, Il17a, Il1f5* (IL-36 receptor antagonist) *and Il1f6* (IL-36alpha) were reduced during resolution (Fig. 2b). In C57BL/6 skin, transcriptional responses were minimal, with only *Mmp12* meeting differential expression criteria (Supplementary Fig. 2c), indicating that large-scale remodeling is restricted to the inflamed tissue context.

KEGG pathway analysis of disease-associated differentially expressed genes revealed enrichment of IL-17 signaling, cytokine-cytokine receptor interactions, and pathways annotated in rheumatoid arthritis and inflammatory bowel disease, reflecting inflammatory circuitry shared across chronic immune-mediated disorders (Fig. 2c, upper; Supplementary Tables 4-5). In parallel, pathways governing actin cytoskeletal organization, calcium signaling, and cellular contractility, frequently annotated as cardiomyopathy or neuronal signaling pathways, were suppressed in disease, consistent with disruption of conserved epithelial structural and signaling programs in inflamed skin (Fig. 2c, lower). During resolution, these patterns shifted: inflammatory and cytokine signaling pathways were attenuated, whereas calcium signaling, MAPK-associated regulation, metabolic pathways, and epithelial signaling programs were re-engaged (Fig. 2d; Supplementary Tables 6-7), indicating coordinated correction of inflammatory and structural gene modules.

Layer-resolved analysis showed that these disease-associated changes, and their reorganization during resolution, were predominantly epidermal. Effect size distributions revealed coordinated shifts across basal, differentiated, and keratinized keratinocyte gene programs, with movement toward baseline expression during resolution (Fig. 2e), indicating that large-scale transcriptional remodeling is executed primarily within keratinocyte compartments.

To further organize these relationships, disease-associated and resolution-responsive genes were grouped into functional modules encompassing inflammatory signaling, keratinocyte stress and differentiation, lipid and fatty acid metabolism, cytoskeletal organization, calcium-dependent signaling, and barrier formation (Fig. 2f). This analysis defines two opposing transcriptional states in chronically inflamed skin: an inflammatory/stress state marked by IL-17 signaling, neutrophil activation, and protease activity, and a homeostatic epidermal state integrating differentiation, barrier integrity, lipid metabolism, calcium signaling, and bioenergetic capacity. Resolution is characterized by suppression of the former and re-engagement of the latter.

Together, these findings position chronic skin inflammation as a keratinocyte-centered transcriptional state in which inflammatory, structural, and metabolic programs are tightly integrated, and show that restoration involves reorganization of this network rather than selective inhibition of individual cytokine pathways.

### Metabolic and calcium-linked pathways are reorganized during resolution

Given enrichment of lipid, bioenergetic, and calcium-associated pathways among genes normalized during resolution (Fig. 2), we examined whether restoration reflects coordinated shifts in metabolic and mechanical gene programs. Directionally normalized genes were grouped into curated metabolic-mechanical axes comprising actin cytoskeleton organization, calcium handling, lipid synthesis and remodeling, cholesterol and sterol metabolism, and fatty acid oxidation and lipid catabolism (Fig. 3a; Supplementary Table 8).

**Figure 3.**
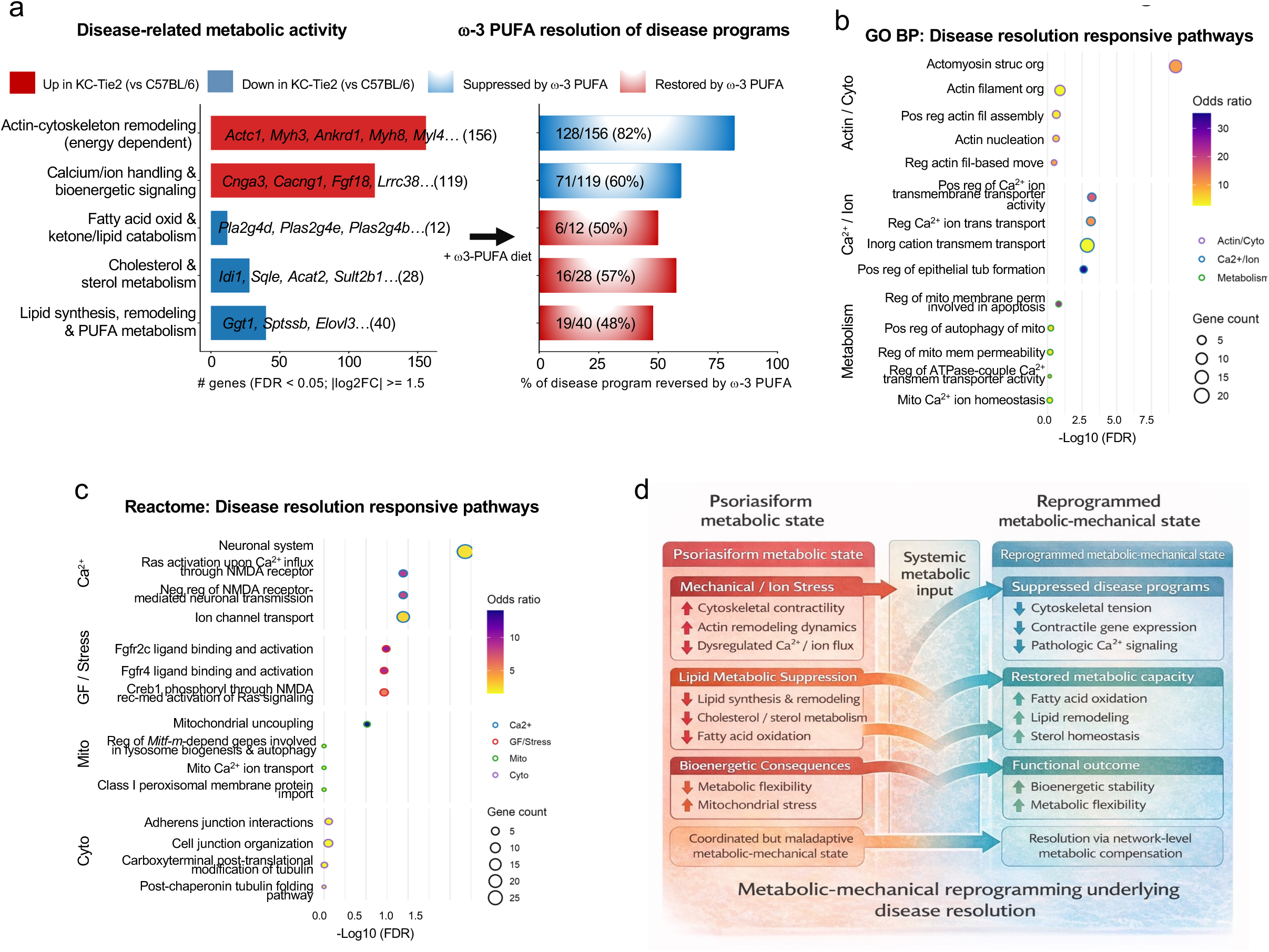
Resolution involves coordinated metabolic and structural pathway shifts. **a,** Disease-associated metabolic gene programs in KC-Tie2 skin and their coordinated, axis-level modulation during disease restoration. Differentially expressed genes in KC-Tie2 skin relative to C57BL/6 controls (FDR < 0.05; |log₂FC| ≥ 1.5) were organized into five curated metabolic-mechanical axes: actin-cytoskeletal remodeling, calcium and ion handling with associated bioenergetic signaling, lipid synthesis and remodeling, cholesterol and sterol metabolism, and fatty-acid oxidation and lipid catabolism. Bars indicate the number of genes increased (red) or decreased (blue) in disease; representative genes are shown, with complete gene-to-axis assignments provided in Supplementary Table 8. The right panel shows the proportion of disease-associated genes within each axis that were directionally modulated in the opposite direction during restoration (FDR < 0.05). **b,** Gene Ontology (GO) Biological Process enrichment of disease-resolution-responsive genes in KC-Tie2 skin, grouped into actin/cytoskeletal remodeling (Actin/Cyto), calcium and ion handling (Ca²⁺/Ion), and metabolic/mitochondrial processes (Metabolism). Dot size reflects gene count, color indicates odds ratio, and the x-axis shows enrichment significance (–log₁₀ FDR). Full GO term statistics and gene lists are provided in Supplementary Table 9. **c,** Reactome pathway enrichment of disease-resolution-responsive genes in KC-Tie2 skin, highlighting coordinated regulation of calcium-linked excitability, growth factor- and stress-responsive signaling, mitochondrial processes, and cytoskeletal/junctional organization. Dot size indicates gene count, color denotes odds ratio, outline color indicates pathway category, and the x-axis shows enrichment significance (–log₁₀ FDR). Full Reactome pathway statistics and gene lists are provided in Supplementary Table 10. **d,** Schematic model integrating transcriptional and pathway-level analyses, illustrating how resolution of psoriasiform inflammation is associated with coordinated reprogramming of metabolic and mechanical gene networks. Disease is characterized by cytoskeletal tension, dysregulated calcium/ion signaling, and suppressed lipid metabolism, whereas resolution involves suppression of disease-associated mechanical and ion-handling programs alongside partial recovery of lipid metabolic capacity, consistent with network-level metabolic compensation rather than global transcriptomic reversal.

This axis-based organization revealed that inflammation in KC-Tie2 skin is characterized by coordinated imbalance across these programs. Disease-associated transcriptional changes included induction of contractile and cytoskeletal genes (*Acta2, Tagln, Myh11, Cnn1*) together with increased calcium-handling components (*Cacna1c, Cacna2d1, Atp2b4*), indicating coordinated activation of structural and calcium-associated regulatory programs within the inflamed epidermis. In contrast, lipid-centered metabolic pathways, including lipid synthesis and remodeling (*Elovl3, Ggt1, Scd1, Dgat2*), cholesterol and sterol metabolism (*Hmgcr, Sqle, Idi1, Fdps*), and fatty acid oxidation (*Pla2g4-*, *Cpt1b, Acadl, Acox1*), were broadly suppressed. During resolution, cytoskeletal and calcium-associated programs elevated in disease were attenuated, whereas lipid metabolic pathways suppressed in KC-Tie2 skin showed coordinated recovery (Fig. 3a). Resolution therefore reflected structured, axis-specific shifts rather than uniform reversal of all disease-associated transcripts.

Independent Gene Ontology Biological Process and Reactome enrichment analyses of genes increased during disease resolution reinforced this organization, identifying coordinated regulation of cytoskeletal organization, calcium transport, epithelial junctional remodeling, and growth factor- and stress-responsive signaling pathways (Fig. 3b,c; Supplementary Tables 9-10). Mitochondrial processes, including ATPase-coupled transport, mitochondrial Ca²⁺ homeostasis, uncoupling, and mitophagy-related pathways, were also enriched, linking bioenergetic adaptation to calcium-associated and lipid metabolic programs rather than indicating an isolated metabolic shift.

These relationships are summarized schematically in Fig. 3d. Resolution was characterized by attenuation of disease-associated cytoskeletal and calcium-linked programs alongside induction of lipid utilization, mitochondrial bioenergetic pathways, and epithelial junctional processes. Together, these findings indicate that inflammatory resolution involves coordinated redistribution of metabolic and structural gene programs rather than uniform reversal of individual transcripts.

### Single-cell analysis localizes inflammatory resolution to keratinocytes

Bulk RNA-seq analyses (Figs. 2-3) revealed coordinated reprogramming of inflammatory, metabolic, and cytoskeletal pathways during resolution of chronic skin inflammation. To determine where these bulk-defined programs are executed at cellular resolution, we performed single-cell RNA sequencing (scRNA-seq) on adjacent skin samples from KC-Tie2 mice under diseased and restored conditions (Fig. 4; Supplementary Table 11).

**Figure 4.**
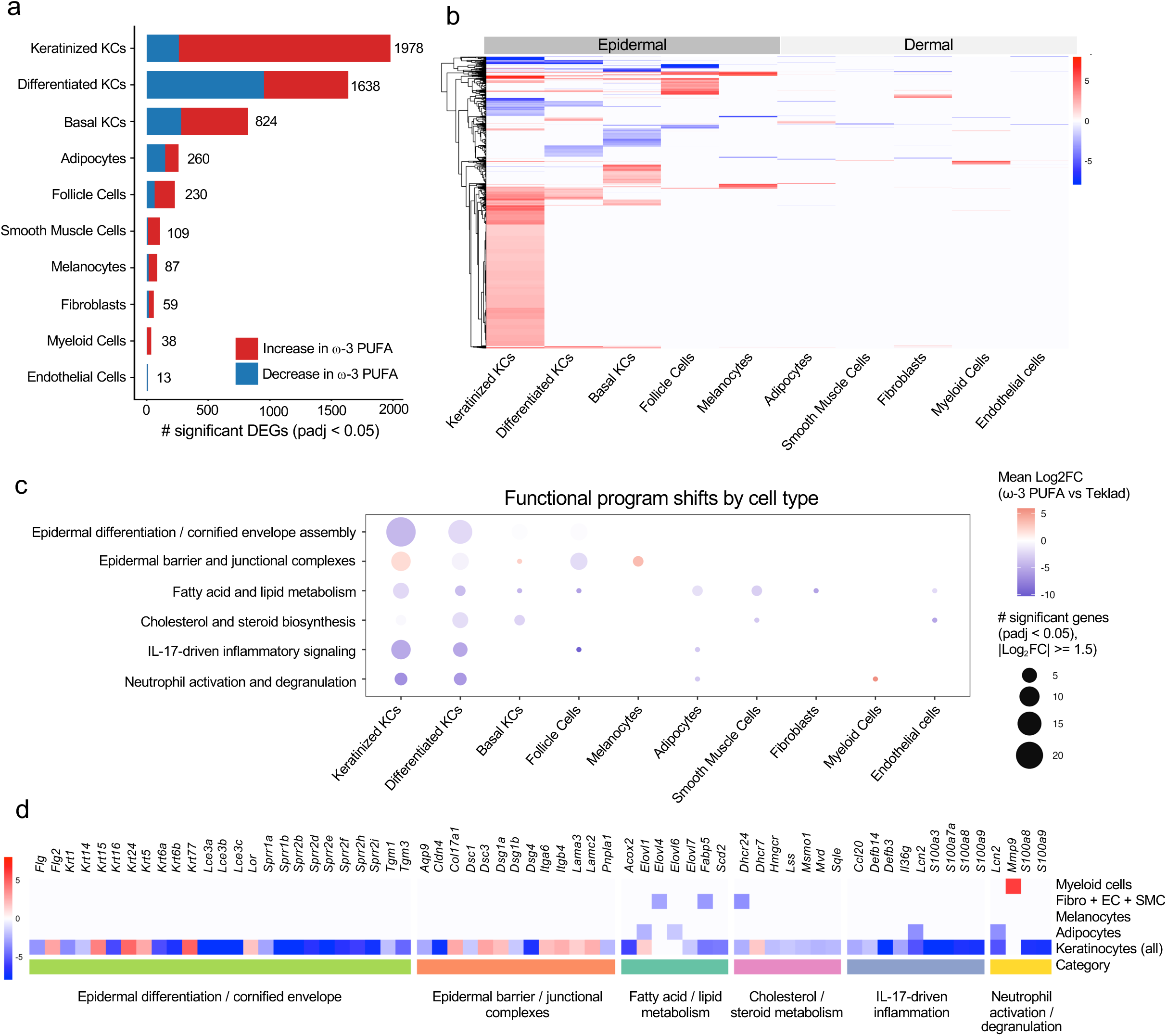
Single-cell analysis localizes transcriptional remodeling to keratinocytes. **a**, Number of significantly differentially expressed genes (DEGs; adjusted *P*C<C0.05) identified by single-cell RNA-seq across major skin cell types in KC-Tie2 skin under restored versus diseased conditions. Bars are colored by direction of regulation, with downregulated genes shown in blue and upregulated genes shown in red. The total number of DEGs per cell type is indicated. Cell-type composition and dataset characteristics are provided in Supplementary Table 11, and the numeric DEG counts underlying this panel are provided in Supplementary Table 13. **b**, Heatmap of a subset of disease resolution-associated DEGs across scRNA-seq–defined skin cell types. Within each cell type, genes were filtered (adjusted *P*C<C0.05; |avg log₂FC|C≥C1.5), and a fixed number of the most strongly induced and suppressed transcripts were selected to visualize dominant transcriptional patterns. Rows represent the union of selected DEGs per cell type and are hierarchically clustered. Columns represent annotated cell types (ordered biologically) and are grouped by epidermal versus dermal compartments. Cell values denote average log₂ fold change (restored vs diseased) for each gene within each cell type; genes not selected in a given cell type are displayed as 0 (white). Complete scRNA-seq DEG statistics are provided in Supplementary Table 12. **c**, Functional program shifts by cell type associated with disease resolution. Bubble plot summarizing the distribution of significant DEGs across predefined biological programs and skin cell types. Dot size represents the number of significantly regulated genes within each program and cell type (adjusted *P*C<C0.05; |avg log₂FC|C≥C1.5). Color indicates the mean log₂ fold change for genes within each category (restored vs diseased), with red denoting induction and blue denoting suppression. Gene-level statistics are provided in Supplementary Table 12, with compartment-stratified pathway enrichment analyses shown in Supplementary Figures 3-5 and detailed in Supplementary Tables 14-25. **d**, Heatmap summarizing representative disease resolution-associated genes across selected functional programs and skin cell compartments in single-cell RNA-seq data. Genes shown represent curated subsets of significantly regulated transcripts contributing to epidermal differentiation and barrier formation, lipid and fatty acid metabolism, cholesterol and sterol biosynthesis, IL-17-driven inflammatory signaling, and neutrophil activation and degranulation. Cell colors represent average log₂ fold change (restored vs diseased) within each cell type, with red indicating induction and blue indicating suppression. Full gene-level statistics are provided in Supplementary Table 12.

Across major epidermal and dermal populations, transcriptional responses during disease resolution were highly asymmetric. Quantification of significantly differentially expressed genes (adjusted P < 0.05) showed that differentiated and keratinized keratinocyte subsets accounted for most resolution-associated transcripts, whereas fibroblasts, endothelial cells, melanocytes, adipocytes, smooth muscle cells, and immune populations exhibited substantially fewer differentially expressed genes (DEGs) (Fig. 4a; Supplementary Tables 12-13). Imposing more stringent thresholds (adjusted P < 0.05; |avg log₂FC| ≥ 1.5) yielded the same pattern: heatmap visualization revealed large, directionally coherent gene clusters within keratinocyte populations, while dermal compartments showed comparatively limited and heterogeneous changes under identical criteria (Fig. 4b; Supplementary Table 12). These single-cell data align with bulk RNA-seq analyses (Fig. 2e), localizing both disease-associated dysregulation and its normalization primarily to epidermal programs.

Integration of cell-type distribution, global expression patterns, and program-level organization (Fig. 4a-d) localized resolution-associated shifts to differentiated and keratinized keratinocytes, spanning structural, metabolic, and inflammatory gene classes. Differentiation and cornification genes (*Flg, Lce* family members, *Tgm1, Tgm3, Sprr* genes) were induced, accompanied by normalization of structural keratins (*Krt1, Krt6a/b, Krt16*) and coordinated regulation of junctional and adhesive components (*Cldn4, Dsc1-3, Dsg1-3, Itga6, Itgb4*), consistent with restoration of epithelial cohesion and control of hyperproliferation.

Concurrently, lipid metabolic programs required for barrier homeostasis were engaged, including fatty acid elongation (*Elovl3, Elovl6, Elovl7, Fabp5*), sterol biosynthesis (*Hmgcr, Sqle, Mvd, Dhcr24*), and lipid oxidation (*Acox2*), while keratinocyte-derived inflammatory mediators (*S100a8, S100a9, Cxcl1, Cxcl2*) were suppressed. Together, these coordinated shifts indicate that epidermal normalization involves simultaneous structural reinforcement, metabolic reactivation, and intrinsic dampening of inflammatory output rather than passive immune redistribution.

To assess whether these transcriptional shifts reflected coordinated functional reorganization, resolution-associated genes were grouped into predefined biological programs encompassing epidermal differentiation and cornified envelope assembly, barrier and junctional organization, lipid and fatty acid metabolism, cholesterol biosynthesis, inflammatory signaling, and neutrophil activation. Keratinocytes exhibited bidirectional shifts across these categories, with induction of differentiation, barrier, and metabolic modules coupled to suppression of inflammatory and neutrophil-associated programs (Fig. 4c). Non-epidermal populations showed more restricted and cell-type-specific responses.

Dermal fibroblasts regulated genes linked to extracellular matrix organization and stromal signaling (*Col12a1, Ltbp2, Pdgfc, Postn, Spry1*), consistent with matrix remodeling accompanying epidermal restoration. Adipocytes selectively modulated metabolic genes (*Hmgcs2, Fabp5, Slc6a6, Txnip*), indicating metabolic adaptation within the dermal niche. Melanocytes regulated stress-responsive, metabolic, and pigmentation-associated genes (*Il31ra, Cacna1d, Bbox1, Dct, Tyr, Tyrp1*), reflecting compartment-specific adaptive responses captured by scRNA-seq (Supplementary Table 13).

Within the immune compartment, keratinocyte-derived neutrophil-associated transcripts were reduced, whereas myeloid cells showed selective induction of *Mmp9* (Fig. 4d), consistent with matrix remodeling and tissue clearance during resolution rather than sustained inflammatory activation. Although pathogenic CD4⁺ and CD8⁺ T cells are a recognized feature of KC-Tie2 inflammation ^13, 14, 16^, T cells were under-represented in this dataset, likely due to technical limitations of archival FFPE tissue. Bulk RNA-seq nevertheless showed robust suppression of *Il17a* and other T cell-derived cytokines during resolution (Figs. 2-3), indicating attenuation of T cell-driven inflammatory circuits.

Compartment-stratified pathway enrichment analyses further refined these patterns. Epidermal genes induced during resolution were enriched for differentiation, cornification, lipid and sterol metabolism, calcium handling, redox regulation, and cytoskeletal organization, whereas suppressed genes mapped to inflammatory and innate immune pathways (Supplementary Figs. 3-5; Supplementary Tables 14-25). Dermal populations showed narrower enrichment profiles, largely centered on extracellular matrix organization and stromal remodeling.

Together, these analyses resolve tissue-level transcriptional shifts into defined cellular programs, indicating that inflammatory resolution reflects coordinated, cell-type-specific reorganization across the skin microenvironment.

### A triad of epithelial regulators anchors disease resolution

Integration of bulk RNA-seq with a previously published label-free proteomic dataset from KC-Tie2 skin ^15^ revealed concordant disease-associated fold changes at RNA and protein levels (Fig. 5a), indicating that core features of the inflammatory state are preserved across molecular layers. Within this shared transcript–protein space, Serpinb3b, Slc25a5, and Cacna2d1 emerged as consistently regulated in disease at both levels. *Serpinb3b* and *Slc25a5* were elevated in inflamed skin and reduced during resolution, whereas *Cacna2d1* exhibited the reciprocal pattern, with suppressed expression in disease and restoration during normalization (Fig. 5b; Supplementary Fig. 6a).

**Figure 5.**
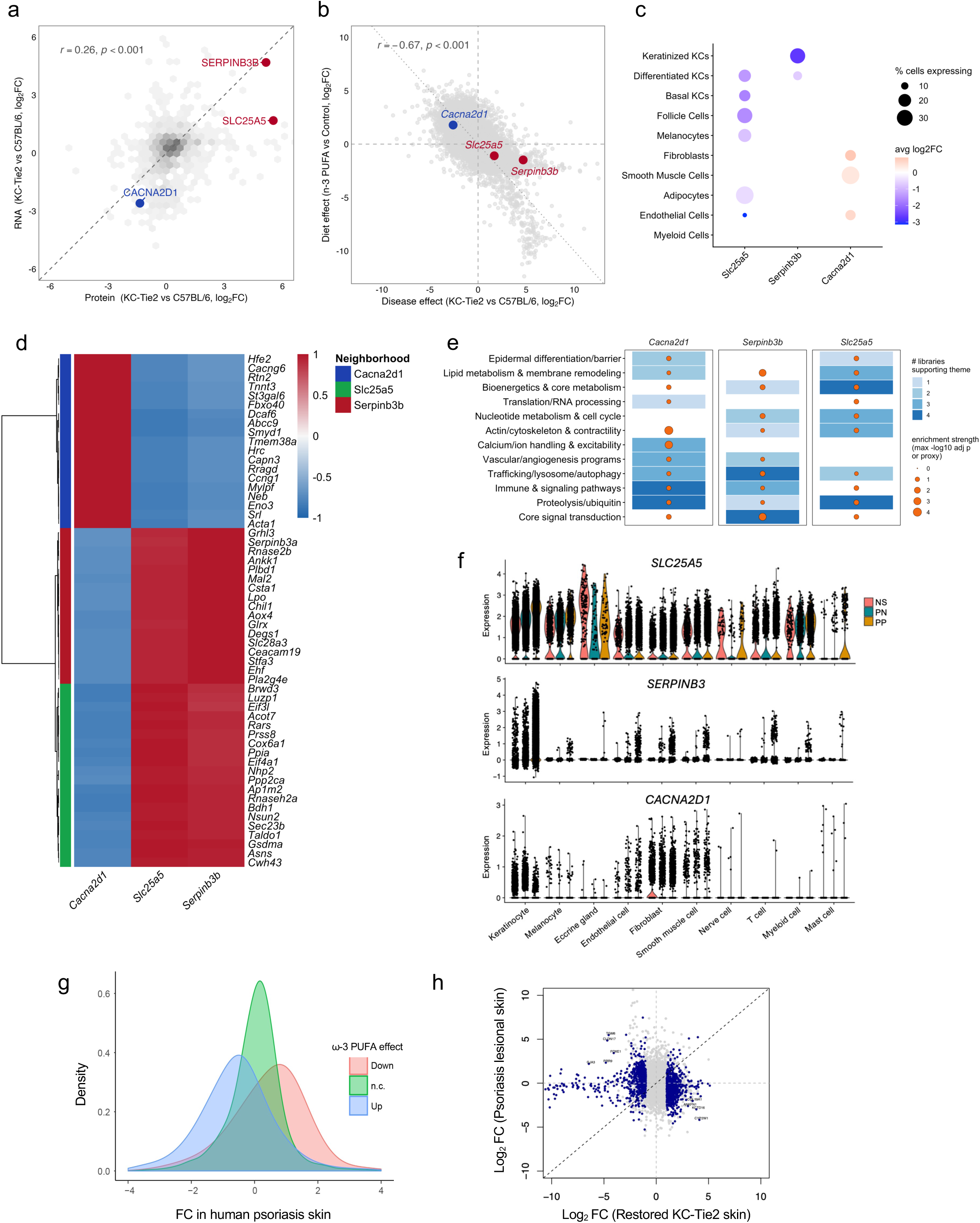
A triad of epithelial regulators anchors disease resolution. **a,** Concordance between transcriptomic and proteomic changes in KC-Tie2 skin. Log₂ fold changes for genes detected in both bulk RNA-seq and previously published label-free proteomics datasets show significant correlation, with Serpinb3b, Slc25a5, and Cacna2d1 highlighted. **b,** Positioning of triad genes within the disease-resolution landscape. Disease effects (KC-Tie2 vs C57BL/6) are plotted against resolution effects within KC-Tie2 skin (ω-3 PUFA vs control). *Serpinb3b* and *Slc25a5 l*ocalize to the disease-elevated/resolution-suppressed quadrant, whereas *Cacna2d1* shows the reciprocal pattern, consistent with repression in disease and restoration during resolution. **c,** Single-cell RNA-sequencing-based cellular distribution of triad gene expression across skin populations. Dot size represents the percentage of cells expressing each gene, and color indicates average log₂ expression. **d,** Triad-centered transcriptional neighborhoods. Heatmap shows Pearson correlations between each triad gene and its 20 most strongly co-varying transcripts across bulk RNA-seq samples, revealing distinct gene-centered neighborhoods (Supplementary Table 26). **e,** Enrichment themes associated with triad neighborhoods across multiple pathway libraries. Tile color indicates the number of independent databases (Gene Ontology Biological Process, KEGG, Reactome, WikiPathways) supporting each biological theme (Supplementary Tables 27-28). **f,** Human single-cell RNA-seq validation of triad gene expression in chronic inflammatory skin disease. Violin plots show expression of *SLC25A5, SERPINB3,* and *CACNA2D1* in keratinocyte and stromal populations from healthy (NS), nonlesional (PN), and lesional (PP) psoriasis skin. *SLC25A5* and *SERPINB3* are increased in lesional epidermis, whereas *CACNA2D1* is reduced. **g,** Distribution of human psoriasis lesion fold changes for genes stratified by direction of regulation during KC-Tie2 resolution. Density plots show that genes suppressed during resolution in KC-Tie2 skin are enriched among transcripts elevated in psoriatic lesions, and vice versa. **h,** Genome-wide comparison of transcriptional effect sizes between restored KC-Tie2 skin and human psoriasis lesions. Opposing directionality across the transcriptome indicates alignment between experimental resolution and human disease-associated expression patterns.

Single-cell RNA sequencing provided cellular context for the triad. Serpinb3b and Slc25a5 were enriched in keratinocyte populations, whereas Cacna2d1 transcripts were less frequently detected in keratinocytes and were more apparent in stromal and vascular compartments (Fig. 2e, Fig. 5c; Supplementary Fig. 6b). Reduced keratinocyte detection of Cacna2d1 likely reflects under-sampling of low-abundance calcium channel transcripts in archived tissue rather than true absence, as CACNA2D1 is readily detectable in healthy keratinocytes (Supplementary Fig. 7). These findings place the triad within partially overlapping but distinct cellular contexts.

To define the transcriptional programs associated with each gene, we performed an unbiased co-variation (“neighborhood”) analysis across the full bulk RNA-seq dataset restricted to protein-coding genes (Supplementary Tables 1-2). For each triad member, Pearson correlation coefficients were calculated against all transcripts, and the 20 most strongly correlated genes were assembled into gene-centered neighborhoods (Fig. 5d; Supplementary Table 26).

Despite minimal overlap in composition, each neighborhood exhibited coherent and biologically distinct patterns of co-regulation (Fig. 5d). The *Cacna2d1*-centered neighborhood was enriched for genes involved in calcium handling, actin cytoskeletal organization, focal adhesion signaling, and contractile programs, including *Kcnj8, Atp1b1, Acta2, Tagln*, and *Myh11*. In contrast, the *Slc25a5* neighborhood was dominated by genes supporting bioenergetic and biosynthetic capacity, including *Nme1, Nme2, Rrm2, Psma5,* and *Eif3e*. The *Serpinb3b* neighborhood was enriched for epithelial stress and lipid remodeling programs, including *Fabp5, Elovl6, Krt16*, *Krt17*, and *S100a10*.

Pathway enrichment of each triad-centered neighborhood identified shared higher-order biological themes, including lipid and membrane metabolism, cytoskeletal organization and mechanotransduction, inflammatory and cytokine signaling, and central metabolic regulation (Fig. 5e; Supplementary Tables 27-28). Lipid-related pathways encompassed fatty acid biosynthesis and oxidation, glycerophospholipid and sphingolipid metabolism, and arachidonic and linoleic acid pathways. Inflammatory modules included IL-1, IL-6, JAK-STAT, MAPK, NF-κB, and Ras signaling, indicating engagement of multiple cytokine-responsive networks across neighborhoods.

To evaluate translational relevance, we examined *SLC25A5, SERPINB3*, and *CACNA2D1* expression in healthy skin and in nonlesional and lesional samples from patients with psoriasis (Fig. 5f) and atopic dermatitis (Supplementary Fig. 7). Consistent with observations in KC-Tie2 skin, lesional epidermis showed increased *SLC25A5* and *SERPINB3* and reduced *CACNA2D1* across overlapping cell populations relative to nonlesional and healthy skin. Genes suppressed in restored KC-Tie2 skin were enriched among transcripts elevated in human psoriatic lesions, whereas genes induced during resolution corresponded to transcripts reduced in lesional skin (Fig. 5g). Genome-wide effect-size comparisons further demonstrated that transcriptional shifts during KC-Tie2 resolution opposed both the direction and magnitude of psoriasis-associated gene regulation (Fig. 5h). Beyond the triad, cross-species analysis identified additional directionally conserved transcripts, including *TGM6, PRR9, FOXE1*, and *CLDN17* (elevated in inflamed skin and reduced with intervention), and *COL19A1, ANKFN1*, and *CYP2W1* (decreased in disease and restored during resolution).

Together, these data indicate that disease resolution concentrates on a limited set of regulatory programs that are directionally conserved across species and across chronic inflammatory skin diseases.

### Functional interference reveals distinct roles of triad genes

To test whether *Cacna2d1, Serpinb3b*, and *Slc25a5* actively modulate disease severity, we performed in vivo siRNA-mediated silencing of each gene in KC-Tie2 skin. Gene-specific siRNAs were delivered topically to ear skin using a validated approach ^13, 18^, and tissue was collected 14 days after treatment initiation (Fig. 6a). Efficient transcript-level knockdown was confirmed for each target (Supplementary Fig. 8).

**Figure 6.**
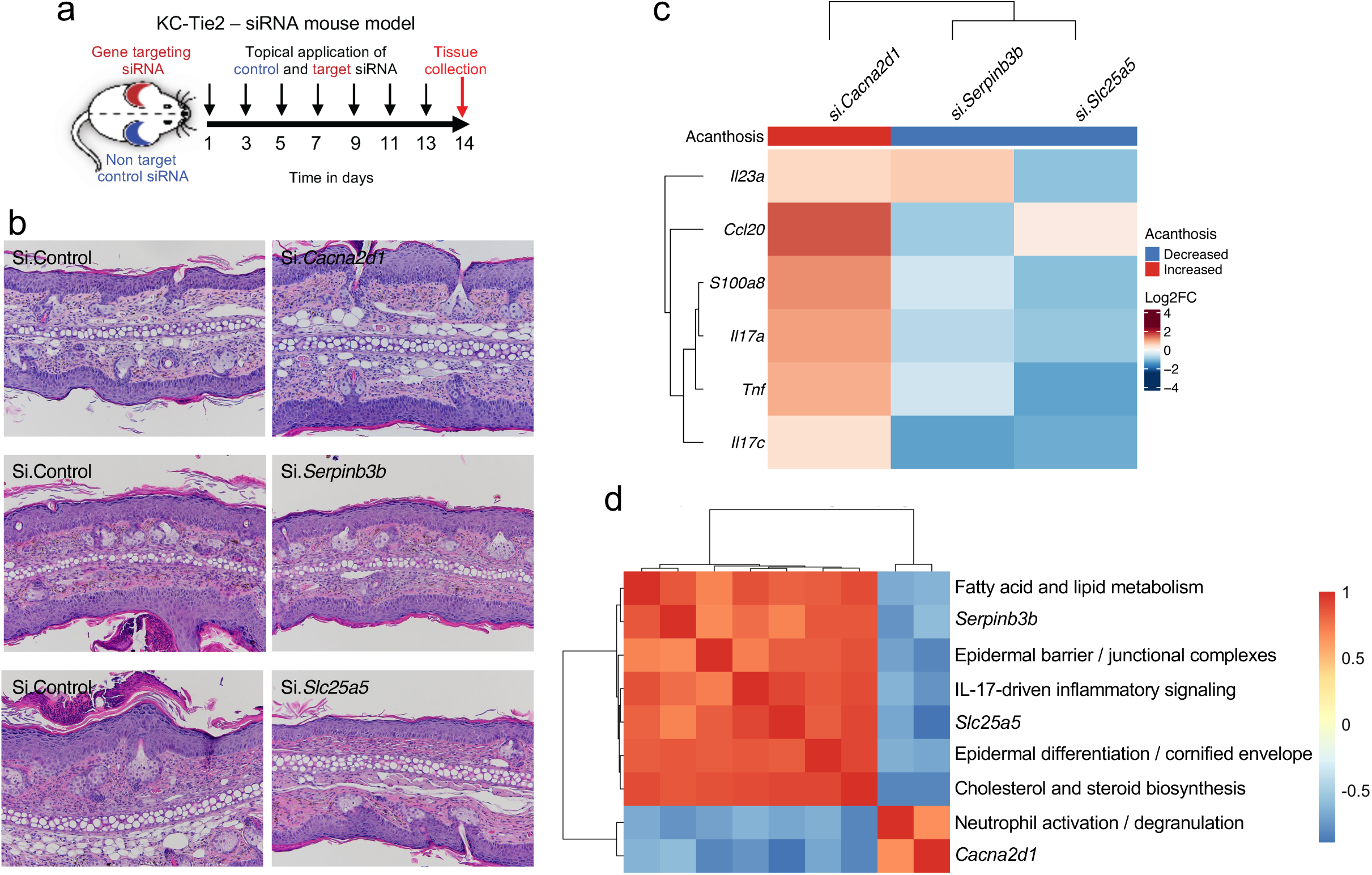
Triad genes exert distinct in vivo control of inflammation. **a,** Experimental schematic for topical siRNA-mediated gene silencing in KC-Tie2 mice. Non-targeting control siRNA or gene-specific siRNAs targeting *Cacna2d1*, *Serpinb3b*, or *Slc25a5* were applied topically to ear skin at the indicated time points, with tissue collection on day 14. **b,** Representative hematoxylin and eosin–stained sections of ear skin following treatment with control siRNA or gene-specific siRNAs. **c,** Heatmap summarizing changes in epidermal acanthosis and expression of IL-23/IL-17-axis-associated genes (*Il23a*, *Ccl20*, *S100a8*, *Il17a*, *Tnf*, *Il17c*) following gene-specific knockdown. Values are shown as log₂ fold change relative to control siRNA-treated skin. Acanthosis is coded as increased or decreased relative to control. **d,** Correlation heatmap relating each targeted gene to transcriptional programs identified by bulk RNA-seq pathway analyses, including fatty acid and lipid metabolism, epidermal barrier and junctional organization, IL-17-associated inflammatory signaling, epidermal differentiation and cornified envelope formation, cholesterol and sterol biosynthesis, and neutrophil activation/degranulation. Quantitative log₂ fold-change values underlying the cytokine and alarmin heatmap in **c** are provided in Supplementary Table 29.

Silencing *Cacna2d1,* which is suppressed in disease and restored during resolution, exacerbated epidermal pathology. KC-Tie2 mice receiving *Cacna2d1* siRNA developed increased acanthosis, parakeratosis, and dermal inflammatory infiltration compared with non-targeting controls (Fig. 6b; Supplementary Fig. 9a). This was accompanied by elevated expression of *Il23a, Il17a, Il17c, Ccl20*, and *S100a8,* indicating amplification of IL-23/IL-17-associated and alarmin-linked inflammatory programs (Fig. 6c; Supplementary Table 29).

In contrast, knockdown of *Serpinb3b* attenuated disease features. Epidermal thickness and inflammatory cell infiltration were reduced, alongside decreased expression of keratinocyte-derived mediators including *Ccl20* and *Il17c* (Fig. 6b-c; Supplementary Fig. 9b).

Targeting *Slc25a5* produced a distinct inflammatory profile. *Slc25a5* silencing significantly reduced *Il23a* and *Il17a* expression, with comparatively modest effects on chemokines such as *Ccl20* (Fig. 6c; Supplementary Fig. 9c), consistent with selective modulation of cytokine output rather than broad suppression of inflammatory signaling.

Program-level correlation analysis further separated these perturbations (Fig. 6d; Supplementary Table 30). *Serpinb3b* and *Slc25a5* associated predominantly with epithelial differentiation and metabolic modules, whereas *Cacna2d1* aligned with inflammatory and neutrophil-related pathways, indicating that each gene regulates a distinct transcriptional domain.

Together, these in vivo perturbations establish that the triad comprises mechanistically active regulators with separable functions in epidermal inflammation. *Cacna2d1* restrains inflammatory amplification, *Serpinb3b* supports epithelial-immune signaling, and *Slc25a5* modulates cytokine output through metabolic control, demonstrating that disease resolution engages multiple regulatory axes rather than a single linear pathway.

### Network architecture of disease resolution

To integrate gene-level and functional findings across preceding analyses, we examined the global organization of disease-associated transcripts that reverse during tissue restoration. Pairwise gene-gene correlations were calculated across bulk RNA-seq samples from KC-Tie2 and C57BL/6 skin using the disease-associated gene set defined in Fig. 2a (Fig. 7; Supplementary Table 2).

**Figure 7.**
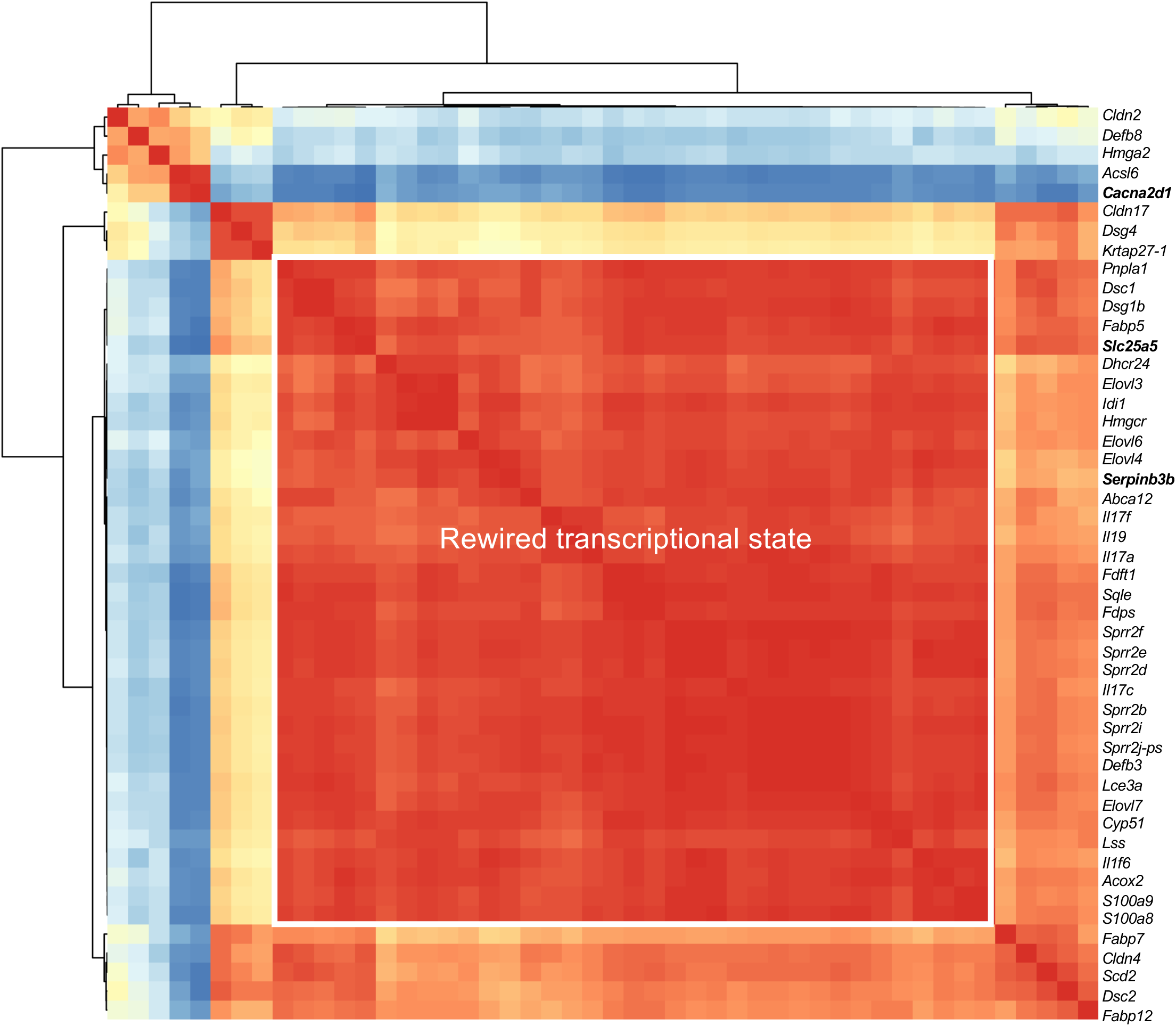
Correlation structure of disease- and resolution-associated gene networks. Hierarchical clustering of pairwise Pearson correlation coefficients calculated across bulk RNA-seq samples using normalized expression values. Each cell represents the correlation between expression profiles of two genes across all samples. Genes were selected from transcripts dysregulated in KC-Tie2 skin and directionally reversed during disease resolution and include markers of epidermal differentiation and cornified envelope formation (e.g., *Krt*, *Sprr*, *Lce* family members), barrier and junctional components, lipid and cholesterol metabolism (e.g., *Elovl*, *Scd*, *Hmgcr*), antimicrobial and IL-17-responsive inflammatory genes (e.g., *Defb*, *Il17a*, *Il17f*, *Il17c*), and neutrophil-associated transcripts. Genes are ordered by hierarchical clustering based on correlation similarity. The boxed region highlights a subset of genes exhibiting high mutual correlation across samples. The disease resolution-associated genes *Serpinb3b*, *Slc25a5*, and *Cacna2d1* are indicated. Color scale denotes Pearson correlation coefficients.

Hierarchical clustering of Pearson correlation coefficients revealed a dominant, densely interconnected transcriptional module spanning epidermal differentiation and cornified envelope genes, lipid and cholesterol metabolic pathways, barrier components, and IL-17-responsive inflammatory mediators. Rather than segregating into discrete pathways, these programs formed a contiguous network, indicating that metabolic, structural, and inflammatory features of disease are embedded within a shared transcriptional architecture.

Neutrophil-associated transcripts formed a partially coupled but separable cluster, suggesting that myeloid programs track with epidermal pathology yet remain peripheral to the central epithelial network.

Within this architecture, *Serpinb3b* and *Slc25a5* localized to the core epidermal module, consistent with their integration into differentiation, metabolic, and inflammatory programs. In contrast, *Cacna2d1* displayed counter-directional correlations relative to the dominant disease block, positioning it as a regulatory node that opposes the prevailing transcriptional configuration.

Together, these analyses define resolution as a shift in network structure rather than selective cytokine attenuation. Chronic skin inflammation is maintained by a highly integrated epidermal transcriptional architecture, and restoration reflects its organized reconfiguration.

## Discussion

Chronic inflammatory skin diseases such as psoriasis and atopic dermatitis have long been conceptualized as immune-driven disorders, a view reinforced by the clinical success of IL-23/IL-17 blockade in psoriasis and IL-4/IL-13 inhibition in atopic dermatitis ^19, 20, 21, 22^. Although immune pathways are required for disease expression, this framework implicitly positions epidermal keratinocytes as downstream responders to immune dysregulation. Our data support a revised hierarchy. Across transcriptomic, cross-species, and functional perturbation analyses, restoration of keratinocyte-intrinsic homeostasis was sufficient to destabilize pathogenic inflammatory circuits in chronically inflamed skin. Keratinocytes therefore emerge not as passive targets, but as regulators of inflammatory persistence whose intrinsic metabolic and structural state determines whether inflammation is sustained or resolved.

Dietary ω-3 PUFA supplementation served as a system-level perturbation that exposed this regulatory logic. In the KC-Tie2 model, intervention reversed thousands of disease-associated transcripts and re-engaged programs governing epidermal differentiation, lipid metabolism, mitochondrial function, and cytoskeletal organization while attenuating IL-23/IL-17-associated inflammatory networks. Importantly, this normalization occurred without direct immune targeting, indicating that restoration of epithelial equilibrium is sufficient to collapse self-sustaining inflammatory loops.

Single-cell RNA sequencing localized execution of these resolution-associated programs primarily to differentiated and keratinized keratinocytes, which accounted for the majority of transcriptional change. Dermal fibroblasts, adipocytes, melanocytes, endothelial cells, and immune populations exhibited more restricted responses. Pathway-level analyses reinforced this hierarchy: epidermal genes induced during resolution aligned with differentiation, lipid and sterol metabolism, calcium handling, and redox regulation, whereas suppressed genes mapped to IL-17 signaling and innate immune amplification. Dermal populations were enriched largely for extracellular matrix organization and stromal remodeling, consistent with adaptive tissue responses downstream of epidermal stabilization. These findings extend emerging models in which keratinocytes actively shape immune homeostasis ^2, 3, 17, 23^, by demonstrating that epithelial state is not merely permissive for resolution but mechanistically sufficient to destabilize chronic inflammatory circuits.

Non-epidermal compartments nevertheless participated in a coordinated multicellular resolution state. Adipocytes exhibited metabolic adaptation, fibroblasts and melanocytes engaged extracellular matrix and stress-associated programs, and myeloid populations showed matrix-remodeling signatures despite suppression of neutrophil-associated transcripts within keratinocytes. These aligned but secondary responses suggest that epidermal recalibration provides the dominant organizing influence, with stromal and immune adaptations reinforcing tissue repair.

Within this context, cross-platform integration of transcriptomic and proteomic data identified a compact triad of regulators, *Serpinb3b, Slc25a5*, and *Cacna2d1,* linking epithelial stress signaling, metabolic state, and calcium-dependent control. Concordant regulation of orthologous genes in human psoriasis and atopic dermatitis supports conservation of this regulatory architecture across species. Functional intervention established that these genes are mechanistically active rather than correlative markers: *Serpinb3b* silencing reduced keratinocyte-derived inflammatory mediators, *Slc25a5* knockdown selectively dampened IL-23/IL-17A output, and loss of *Cacna2d1* exacerbated epidermal pathology and inflammatory amplification. Together, these divergent phenotypes indicate that inflammatory resolution is governed by parallel epithelial control nodes integrating stress responses, bioenergetic capacity, and cytokine regulation.

Resolution, therefore, reflects reconfiguration of network topology rather than isolated suppression of cytokine expression. This systems-level organization provides a framework for understanding why cytokine-directed therapies can effectively attenuate disease activity, yet may not uniformly re-establish durable tissue homeostasis, whereas epithelial reprogramming disrupts the broader transcriptional architecture sustaining inflammation.

The implications extend beyond the skin. Barrier epithelia in the gut, lung, and other mucosal tissues must coordinate metabolic activity, structural integrity, and immune surveillance to maintain homeostasis. Disruption of epithelial regulatory programs in these tissues is increasingly recognized as a driver of chronic inflammatory disease ^24, 25, 26, 27^. Our findings suggest that epithelial state itself functions as an organizing principle of inflammatory persistence or resolution. Rather than serving solely as targets of immune dysregulation, epithelial cells may act as central regulatory hubs whose metabolic and structural programs determine the stability of tissue-level immune circuits across barrier organs,

## Methods

### Study approval

Human samples were obtained from volunteer patients with chronic inflammatory skin disease and healthy controls with informed written consent before inclusion in the study in accordance with Declaration of Helsinki principles. All protocols were approved by the University of Michigan institutional review board. All animal experiments were approved by the Case Western Reserve University institutional animal care and use committee and conformed to the American Association for Accreditation of Laboratory Animal Care guidelines.

### Sex as a biological variable

Males and females were used in the human and animal experiments.

### Mouse experiments

KC-Tie2 mice on a C57BL/6 background were generated as previously described ^13, 28^. Briefly, K5tTA mice ^29^ were mated with TetosTie2(*Tek*) ^30^ mice in the presence of doxycycline chow (200mg/kg, Bio-Serve #S3888) for 7 days and then food was replaced with standard P3000 chow (Prolab® IsoPro® RMH 3000; LabDiets, Richmond, Indiana).

Genotyping was completed by Transnetyx Inc. Mice inheriting copies of both genes express Tie2 in K5+ cells (KC-Tie2) that we previously demonstrated to be keratinocyte-specific ^31^. Littermates inheriting neither transgene served as controls.

### Diet manipulation

At 6 weeks of age, cohoused KC-Tie2 and C57BL/6 littermates were switched from P3000 chow to 2018 Teklad chow (Inotiv) or Teklad chow enriched with 10% fish oil (Inotiv). Mice were euthanized at 12 weeks-of-age and matched dorsal skin was collected for histology, immunohistochemistry, and RNA analyses.

### Topical si.RNA application to ear skin of KC-Tie2 mice

ON-TARGETplus siRNA SMARTpool for mouse *Cacna2d1*(gene id: 12293; L-044128), *Serpinb3b* (gene id: 383548; L-051544), *Slc25a5* (gene id: 11740; L-042392) and non-targeting pool (control siRNA, D-001910) were purchased from Dharmacon. A preparation for topical application was made that included 2.5nM siRNA/2µl of Lipofectamine 3000 (Invitrogen) /10mg of over-the-counter generic hand cream. Topical siRNA targeting *Cacna2d1, Serpinb3b, Slc25a5* or control siRNA was applied to ear skin of KC-Tie2 mice (Figure 6a) every other day for 14 days as we have previously described ^13^. This approach enables control siRNA and targeted siRNA to be applied onto the skin of the same mouse, allowing for a paired statistical analysis and eliminating potential confounding factors that could elicit changes, including microenvironment, cage, and littermate variability. Mice were euthanized the day after the last application and skin was collected for histology and RNA analyses.

### Tissue collection and histological and morphometric analyses

After mice were euthanized, hair was shaved and skin from the back was harvested and processed for either frozen or paraffin sectioning. For paraffin sectioning, skin was placed in 10% buffered formalin (Surgipath Medical Industries, Richmond, IL), overnight at 4°C, followed by dehydration and embedding (Sakura Finetech, Torrance, CA). For frozen sectioning, skin was immediately embedded in Tissue Fixation Media in TBS (TFM; Triangle Biomedical) and then flash frozen in liquid nitrogen. Additional pieces of back skin from the same animal were flash frozen in RNA later in liquid nitrogen and stored at -80°C for use in RNA experiments.

H&E staining on skin from KC-Tie2 mice was completed on 5μm thick paraffin sections using standard protocols. Images were captured using a Leica DM L82 microscope with an Olympus SC180 camera and Olympus cellSansEntry (V2.3) software. Epidermal thickness measures were completed using Adobe Photoshop (version 24.0.0). For each animal, ∼10 measurements were taken from at least five different fields of view from one section (∼50 measurements per animal). Epidermal thickness was measured from the stratum basale to stratum granulosum, excluding the stratum corneum. Immunohistochemistry against T cells and myeloid cells was performed on TFM-embedded frozen skin sectioned at 8μm, using antibodies targeting CD4 (Cat# 550280, BD Biosciences), CD8a (Cat# 550281, BD Biosciences), F4/80 (cat# 14-4801-82 EBiosciences) or CD11c (14-0114-82, EBiosciences) following established protocols in our lab ^13, 16, 32, 33^. Antibodies were detected using either rabbit anti-rat IgG biotinylated (CD4, F4/80; Vector Labs, Burlingame, CA) or goat anti-hamster IgG biotinylated (CD11c; Jackson Immunoresearch Labs, West Grove, PA), amplified with Avidin/Biotinylated Enzyme Complex (Vector Labs) and visualized using the enzyme substrate diaminobenzidine (Vector Labs). The slides were counterstained with hematoxylin. Images were captured as above.

### RNA isolation and qRT-PCR

Total RNA was isolated using RNeasy plus kit (Cat #74136, Qiagen) and real-time quantitative PCR was performed on a 7900HT Fast Real-time PCR system (Applied Biosystems) with TaqMan Universal PCR Master Mix (Cat #4304437, ThermoFisher Scientific). Mouse primers (ThermoFisher Scientific) used in this study were: *Il23a*, Mm01160011_g1; *Il17a*, Mm00439618_m1; *Il17c,* Mm00521397_m1, *Ccl20*, Mm00444228_m1; *S100a8*, Mm00495696_g1; *Tnf*, Mm00443260_g1, *Cacna2d1*, Mm00486607_m1; *Serpinb3b*, Mm03032256_uH; *Slc25a5*, Mm00846873_g1; and *Gapdh*, Mm99999915_g1. Relative expression was calculated using the 2^-ΔCt method normalized to *Gapdh*.

### Bulk RNA sequencing and analysis

RNA quality was assessed with a 5200 Fragment Analyzer System and Standard Sensitivity RNA kits (Agilent, Santa Clara, CA). Total RNA was normalized to 250ng prior to random hexamer priming and libraries generated by TruSeq Stranded Total RNA - Globin kits (Illumina Inc., San Diego, CA). The resulting libraries were assessed on the Fragment Analyzer (Agilent) with the High Sense Large Fragment kit and quantified using a Qubit 3.0 fluorometer (ThermoFisher Scientific, Carlsbad, CA). Medium depth sequencing, yielding a minimum of 30 million reads per sample, was performed on a NextSeq 550 (Illumina) using a 75-base pair, paired-end run design.

After adapter trimming, sequencing reads were aligned to the mouse reference genome (mm10) using STAR ^34^, and gene-level counts were generated using HTSeq ^35^. Count matrices and sample metadata were imported into DESeq2 ^36^ in R using a negative binomial generalized linear model with size-factor normalization (median-of-ratios) and differential expression analysis. P values were adjusted for multiple testing using the Benjamini-Hochberg false discovery rate (FDR) procedure. Unless otherwise specified, genes were considered differentially expressed at adjusted P < 0.05 and |log₂ fold change| ≥ 1.5. DESeq2-normalized expression values are provided in Supplementary Table 1, and genome-wide differential expression statistics for all detected genes across experimental contrasts are provided in Supplementary Table 2. Volcano plots, density plots, enrichment analyses, and summary visualizations were generated using ggplot2 (v4.0.1); gene labels were positioned using ggrepel where indicated. Principal component analysis was performed on DESeq2-normalized expression values (Supplementary Table 1). Heatmaps of ω−3 PUFA-reversed differentially expressed genes (Supplementary Fig. 2) were generated in GraphPad Prism (v10.6.1) using normalized expression values exported from DESeq2 (Supplementary Table 1) without hierarchical clustering.

Functional enrichment analysis was performed using Enrichr ^37, 38, 39^ on up- and downregulated gene sets separately. Gene Ontology Biological Process, KEGG, Reactome, and WikiPathways libraries were queried as indicated. Terms with FDR-adjusted P < 0.05 were considered significant. Enrichment results were exported and visualized in R. For selected analyses, differentially expressed genes were grouped into curated biological axes based on gene function and pathway annotation (Supplementary Tables 8-10).

### Cross-omics concordance analysis

To evaluate concordance between transcriptomic and proteomic changes in diseased KC-Tie2 skin, genes detected in both bulk RNA-seq and a previously published label-free proteomics dataset from KC-Tie2 skin ^15^ were intersected. For each shared gene, log₂ fold changes for KC-Tie2 versus C57BL/6 were extracted from each dataset and compared using Pearson correlation. Significance was assessed using a two-sided correlation test, and the correlation plot was generated in R.

### Disease-versus-resolution effect mapping

For each protein-coding gene, disease-associated effects were calculated as log₂ fold change (KC-Tie2 vs C57BL/6) under control diet. Resolution-associated effects were calculated as log₂ fold change (KC-Tie2 restoration diet vs KC-Tie2 control diet). Genes were plotted in a two-dimensional effect space (disease effect on the x-axis; resolution effect on the y-axis) to visualize directionality and identify genes increased in disease but suppressed during resolution (and vice versa). Pearson correlation was computed across genes, and plots were generated in R.

### Gene co-variation and correlation network analyses

To define transcriptional neighborhoods centered on *Cacna2d1, Slc25a5*, and *Serpinb3b*, Pearson correlation coefficients were computed between each triad gene and all other genes using normalized bulk RNA-seq expression values across all samples (pairwise complete observations). For each triad gene, transcripts were ranked by absolute correlation (|r|), and the top correlated transcripts were retained to define gene-centered neighborhoods. Correlation P values were calculated from r and sample size using the corresponding t-distribution.

For visualization, the union of selected transcripts was displayed as a correlation heatmap (r, −1 to 1) with rows hierarchically clustered and columns fixed to triad-gene order. Each transcript was assigned to the neighborhood of the triad gene with which it exhibited the strongest absolute correlation. Full correlation coefficients and associated P values are provided in the supplementary tables.

To assess higher-order transcriptional organization, pairwise Pearson correlations were additionally computed among genes within defined disease-associated gene sets across bulk RNA-seq samples. The resulting gene-gene correlation matrix was hierarchically clustered and visualized as a heatmap to evaluate coordinated transcriptional structure.

Neighborhood enrichment and consensus themes. For each triad neighborhood gene set, functional enrichment was performed using Enrichr against Gene Ontology Biological Process, KEGG, Reactome, and WikiPathways libraries. Enrichment results were exported and aggregated in R by grouping related terms into higher-order biological themes. For each theme, the number of libraries supporting the theme and the maximum enrichment strength (e.g., −log₁₀ adjusted P value or proxy score) were summarized for visualization.

### Cross-species directionality comparison

Published human psoriasis bulk RNA-seq datasets (GSE121212) ^40^ were used to perform effect size comparisons between differentially expressed DEGs in human psoriasis and KC-Tie2 skin. Fold change (log₂) was used as effect size for comparison, only gene orthologs were used in the analysis.

Genes were stratified based on their direction of regulation during resolution in KC-Tie2 skin (increased vs decreased in restoration diet relative to control diet, using the study’s DEG thresholds). For each stratum, log₂ fold changes from an external human psoriasis lesional-versus-nonlesional (or lesional-versus-healthy) bulk transcriptomic dataset were extracted. Distributions of human fold changes were visualized as kernel density plots to test whether genes suppressed during murine resolution were enriched among genes elevated in human lesions (and vice versa).

### Genome-wide effect-size concordance

Genome-wide log₂ fold changes from restored KC-Tie2 skin (restoration diet vs control diet within KC-Tie2) were compared with fold changes from human psoriasis lesional skin (lesional vs comparator) across shared orthologous genes. Effect sizes were plotted against each other for all shared genes and Pearson correlation was calculated to quantify global directional concordance.

### ScRNA-Sequencing

For mouse scRNA-seq analyses, FFPE skin adjacent to tissue used for acanthosis and immune cell quantification was profiled using the Chromium Single Cell Gene Expression Flex (Fixed RNA Profiling) workflow (10x Genomics). Single-cell (fixed RNA) libraries were prepared from mouse FFPE skin sections using the Chromium Single Cell Gene Expression Flex reagent kit (10x Genomics) according to the manufacturer’s instructions. Briefly, samples were hybridized with the Chromium Mouse Transcriptome Probe Set v1.0.1 (10x Genomics), followed by single-cell partitioning and barcoding on a Chromium X Series instrument (10x Genomics). Libraries were sequenced on an Illumina platform.

Sequencing data were processed with Cell Ranger multi (v9.0) for demultiplexing, alignment, and generation of gene-by-cell count matrices. Reads were aligned to the mouse reference genome mm39 (refdata-gex-GRCm39-2024-A). Downstream analysis was performed in Seurat (v5.1.0) using the aggregated raw count matrices ^41^. SoupX (v1.6.2) was used to estimate and remove ambient RNA contamination using default parameters ^42^, and scDblFinder (v1.16.0) was used to identify and remove putative doublets using default parameters ^43^. Filtered count matrices were log-normalized and scaled, and principal component analysis was performed. Batch correction was performed using Harmony (v1.0.1) with donor specified as the batch covariate. Cells were embedded using UMAP and clustered using a graph-based approach. Clusters were annotated using a curated panel of canonical marker genes.

Cells with fewer than 200 detected transcripts and/or greater than 25% mitochondrial reads were excluded. Differentially expressed genes were identified using the likelihood ratio test implemented in Seurat (FindMarkers). Significance was defined as Bonferroni-adjusted P < 0.05 with |log2 fold change| > log2(1.2), unless otherwise indicated.

For visualization of regulated genes across cell types (Figure 4), an additional effect-size threshold of |avg log₂FC| ≥ 1.5 was applied. For selected heatmaps, the top up- and downregulated genes per cell type were selected based on average log₂ fold change. A gene-by-cell type matrix of average log₂ fold changes was constructed from the union of selected genes, and genes not selected in a given cell type were displayed as zero for visualization. Heatmaps were generated in R (v4.5.2) using the pheatmap package (v1.0.13). Genes (rows) were hierarchically clustered using Euclidean distance with complete linkage, and columns were ordered according to predefined biological grouping without clustering.

For visualization of triad gene expression across mouse skin cell types (Figure 5c), a per-cell-type summary table was generated from the cleaned single-cell differential expression results containing average log₂ fold change (avg log₂FC) and the proportion of cells expressing each gene. Dot plots were generated in R using ggplot2, with dot size representing the percentage of cells expressing the gene and dot color representing avg log₂FC (midpoint = 0). Cell types were ordered according to predefined biological groupings without clustering.

For program-level summaries, significantly regulated genes were grouped into predefined biological categories based on curated gene lists. For each program and cell type, the number of significant genes and the mean average log₂ fold change were calculated for visualization.

### Human single-cell datasets

Publicly available single-cell RNA-seq datasets (GSE173706 and GSE147424) from healthy skin and from nonlesional and lesional psoriasis (and atopic dermatitis, where indicated) were used to evaluate expression of *SERPINB3, SLC25A5*, and *CACNA2D1* across annotated skin cell populations. Expression was visualized using violin plots stratified by diagnosis group (healthy, nonlesional, lesional) and cell type annotations provided in the original studies. Plots were generated with by the *VlnPlot* function in Seurat.

Statistical tests and graphing statistical analysis were performed using GraphPad Prism 10 (v10.6.1). Data are shown as the mean and standard error of the mean unless otherwise stated. Exact statistical tests, sample sizes, and significance thresholds are provided in figure legends. Two-way ANOVA with Tukey’s post hoc testing or paired Student’s t-tests were used as indicated. Significance thresholds were defined as *P < 0.05, **P < 0.01, ***P < 0.001, ****P < 0.0001.

## Materials availability

All unique/stable reagents generated in this study are available from the corresponding author on completion of a Materials Transfer Agreement.

## Data and code availability

Date used to generate findings of this study are provided in Supplementary Tables 1-30. FASTQ datasets for mouse skin bulk and single cell RNA sequencing have been deposited at NCBI Gene Expression Omnibus and are publicly available as of the date of publication. Human bulk and single cell RNA sequence data are available at GSE121212, GSE147424, and GSE173706.

## Acknowledgments

This work was supported in part by grants from the NIH including: P50 AR070590 to NLW, TSM, and MJC, R01s AR073196, AR084312 to NLW, P30 AR039750 to NLW and TSM, P30 AR075043 to JEG and LCT, and by grants from the National Psoriasis Foundation to NLW and TSM, and to NLW, LCT and JEG. RNA sequencing was performed with support from the Applied Functional Genomics Core at Case Western Reserve University and the AGC Core Facility at University of Michigan. The funders were not involved in the study design, data collection, data analysis, manuscript preparation and/or publication decisions. The content is solely the responsibility of the authors and does not necessarily represent the official views of the NIH.

## Author contributions

NLW designed the research studies, DG, RGS, JF, JW, MM, BR, MJC, LCT, JEG, and NLW conducted the experiments, NLW, DG, RB, BR, MJC, and LCT acquired the data, and DG, RGS, RB, BR, MJC, LCT, JEG and NLW analyzed the data. AS drafted components of the manuscript, and NLW wrote the manuscript. NLW, LCT, MJC, and JEG revised the manuscript. The work was funded by grants to MJC, LCT, JEG, TSM and NLW.

## Materials & Correspondence

Correspondence and material requests should be addressed to

