## Supplementary Fig. for "Restoration of Keratinocyte Homeostasis Drives Resolution of Skin Inflammation"

The authors have declared that no competing interests exist.

### Supplemental Figures

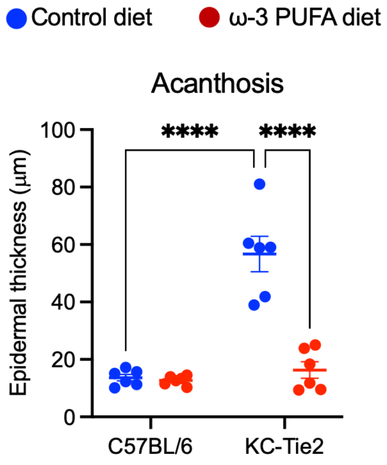


**Supplementary Figure 1. Restoration of epidermal thickness in KC-Tie2 skin.** Quantification of epidermal thickness (acanthosis) in dorsal skin from C57BL/6 and KC-Tie2 mice maintained on either a control diet or a long-chain ω-3 polyunsaturated fatty acid (PUFA)–enriched diet. Each data point represents an individual mouse (n = 6 per group); bars indicate mean ± s.e.m. Statistical significance was assessed by two-way ANOVA with Tukey's multiple-comparisons test. ****$P < 0.0001$.

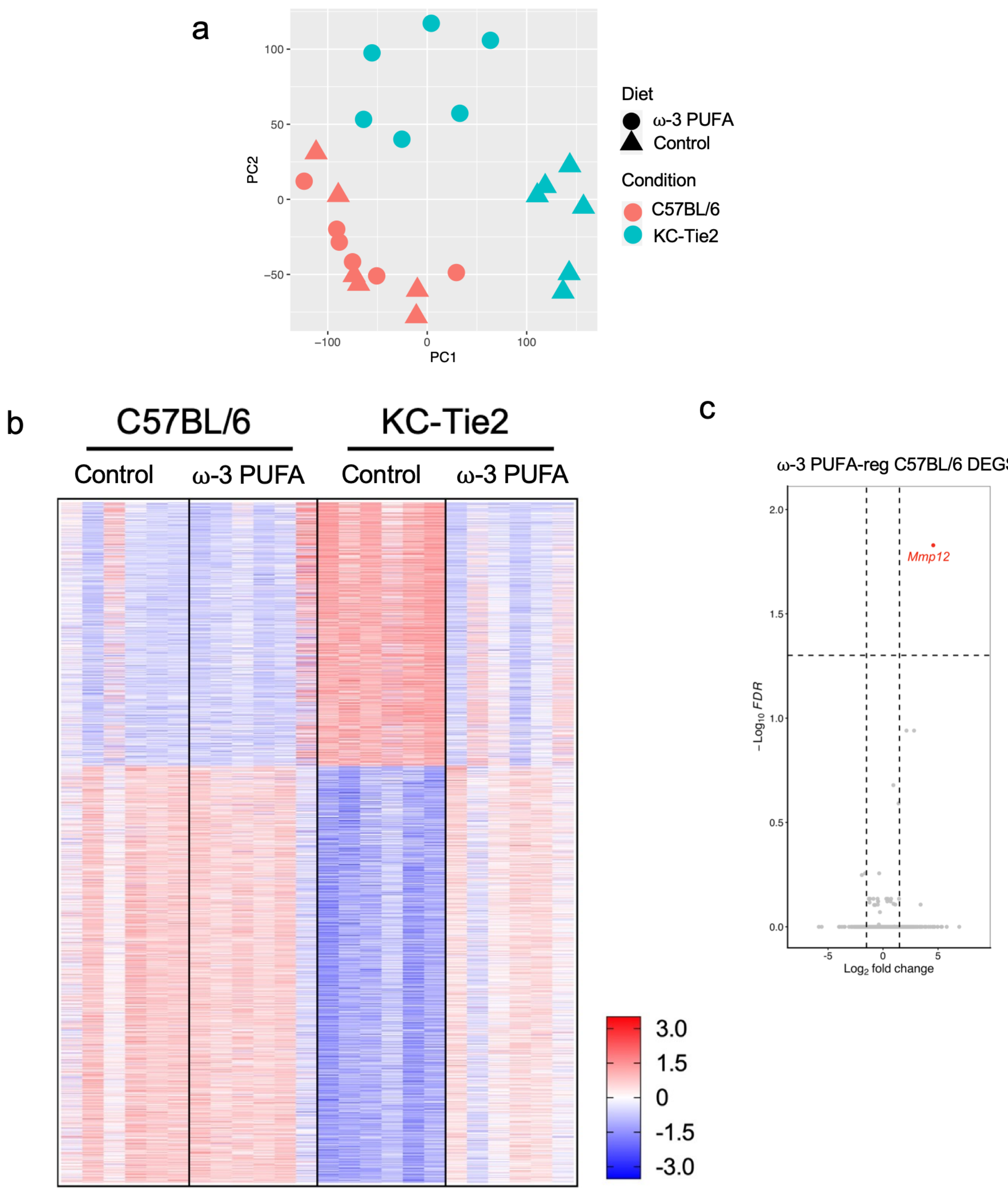


**Supplementary Figure 2. Global transcriptomic structure and disease-context specificity of ω-3 PUFA-responsive gene expression. a,** Principal component analysis (PCA) of bulk RNA-sequencing data from dorsal skin of C57BL/6 and KC-Tie2 mice maintained on either control or ω-3 PUFA-enriched diets (n = 6 per group). Samples are color-coded by genotype and shaped by diet. KC-Tie2 samples segregate strongly from C57BL/6 controls, reflecting a global disease-associated transcriptional state. ω-3 PUFA exposure shifts KC-Tie2 samples toward the wild-type transcriptional space, whereas C57BL/6 samples show minimal separation by diet. **b,** Heatmap of unsupervised hierarchical clustering of DESeq2-normalized gene expression across all samples. Rows represent genes and columns represent individual biological samples grouped by genotype and diet. ω-3 PUFA exposure reverses disease-associated transcriptional patterns in KC-Tie2 skin, restoring expression profiles toward those observed in wild-type controls, while exerting minimal effects in C57BL/6 skin**. c,** Volcano plot of differential gene expression in C57BL/6 skin from ω-3 PUFA-exposed versus control conditions. Genes meeting thresholds of $|\log_2$ fold change$| \geq 1.5$ and adjusted FDR $< 0.05$ are highlighted. Only a single transcript (*Mmp12*) met these criteria, indicating that ω-3 PUFA-responsive transcriptional remodeling is strongly disease-context dependent. Underlying normalized expression values and differential expression statistics are provided in Supplementary Data Tables 1 and 2.

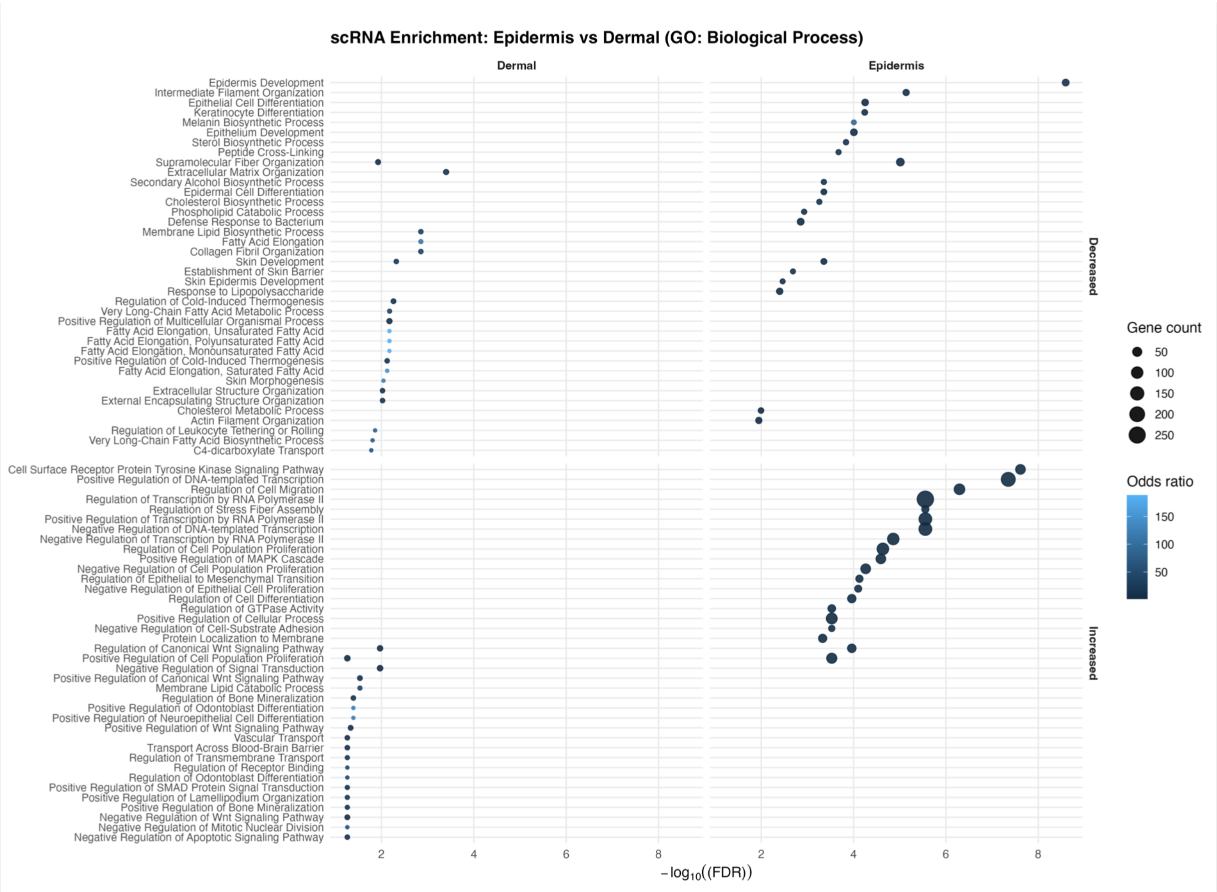


**Supplemental Figure 3. Gene Ontology Biological Process enrichment reveals epidermal-dominant transcriptional programs in single-cell RNA-seq data.** Gene Ontology (GO) Biological Process enrichment analysis of differentially expressed genes identified by single-cell RNA-seq, stratified by epidermal versus dermal skin compartments in KC-Tie2 mice following disease resolution. Enriched GO terms are shown separately for genes decreased (top) or increased (bottom) relative to control conditions. Dot position reflects enrichment significance (-$\log_{10}$ FDR), dot size indicates the number of genes contributing to each term, and color denotes odds ratio. Results highlight preferential enrichment of epidermal differentiation, barrier organization, lipid metabolic, and regulatory processes within epidermal cells, whereas dermal compartments exhibit more limited and distinct functional enrichment patterns. Full GO term statistics and gene lists are provided in Supplementary Tables 14-17.

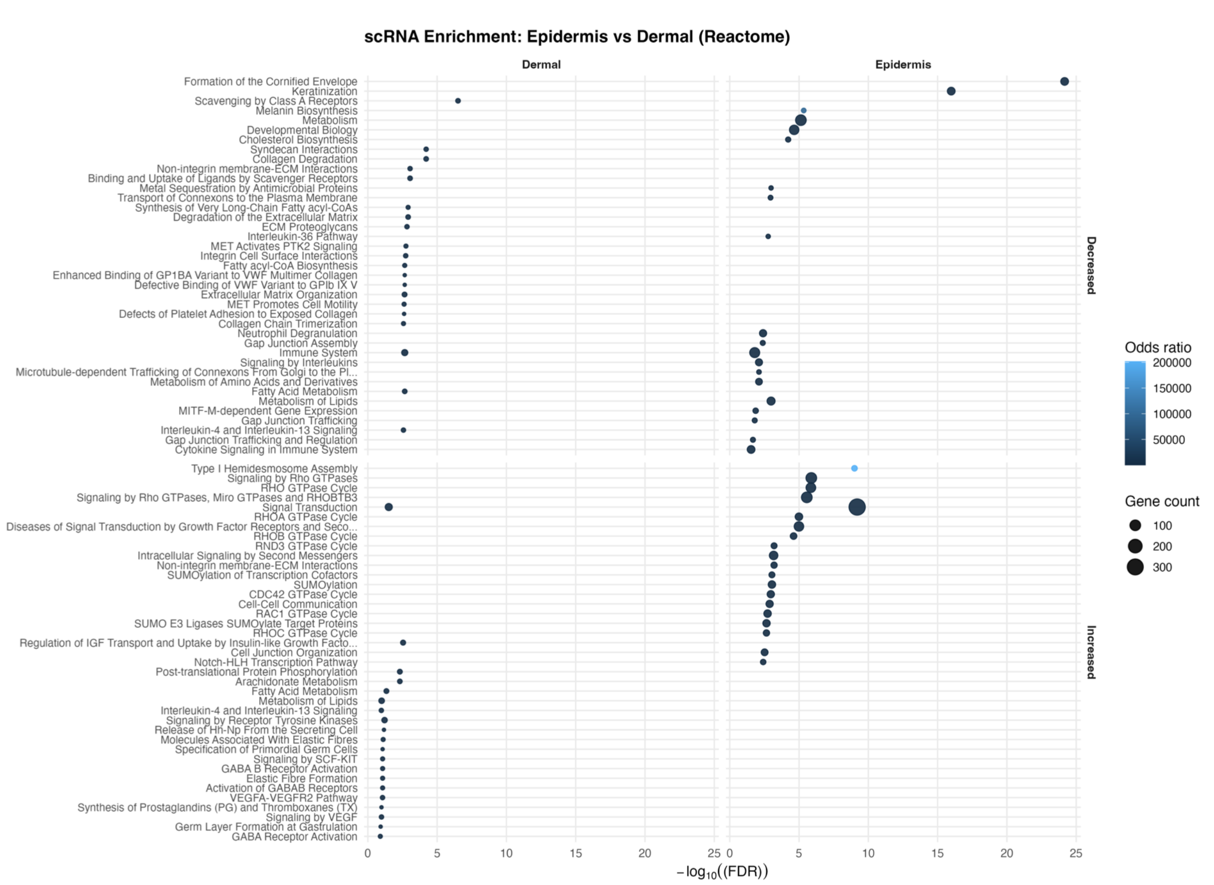


**Supplementary Figure 4. Reactome pathway enrichment highlights epidermal-dominant signaling and metabolic programs during disease resolution in single-cell RNA-seq data.** Reactome pathway enrichment analysis of differentially expressed genes identified by single-cell RNA-seq, stratified by epidermal versus dermal skin compartments in KC-Tie2 mice under restored versus diseased conditions. Enriched Reactome pathways are shown separately for genes decreased (top) or increased (bottom) during disease resolution. Dot position reflects enrichment significance (-$\log_{10}$ FDR), dot size indicates the number of genes contributing to each pathway, and color denotes odds ratio. Epidermal compartments show preferential enrichment of pathways related to keratinization, junctional organization, lipid and fatty acid metabolism, calcium-linked signaling, and stress-responsive processes, consistent with coordinated structural and metabolic reprogramming within keratinocytes. Dermal compartments exhibit more limited and distinct pathway enrichment profiles. Full Reactome pathway statistics and gene lists are provided in Supplementary Tables 18-21.

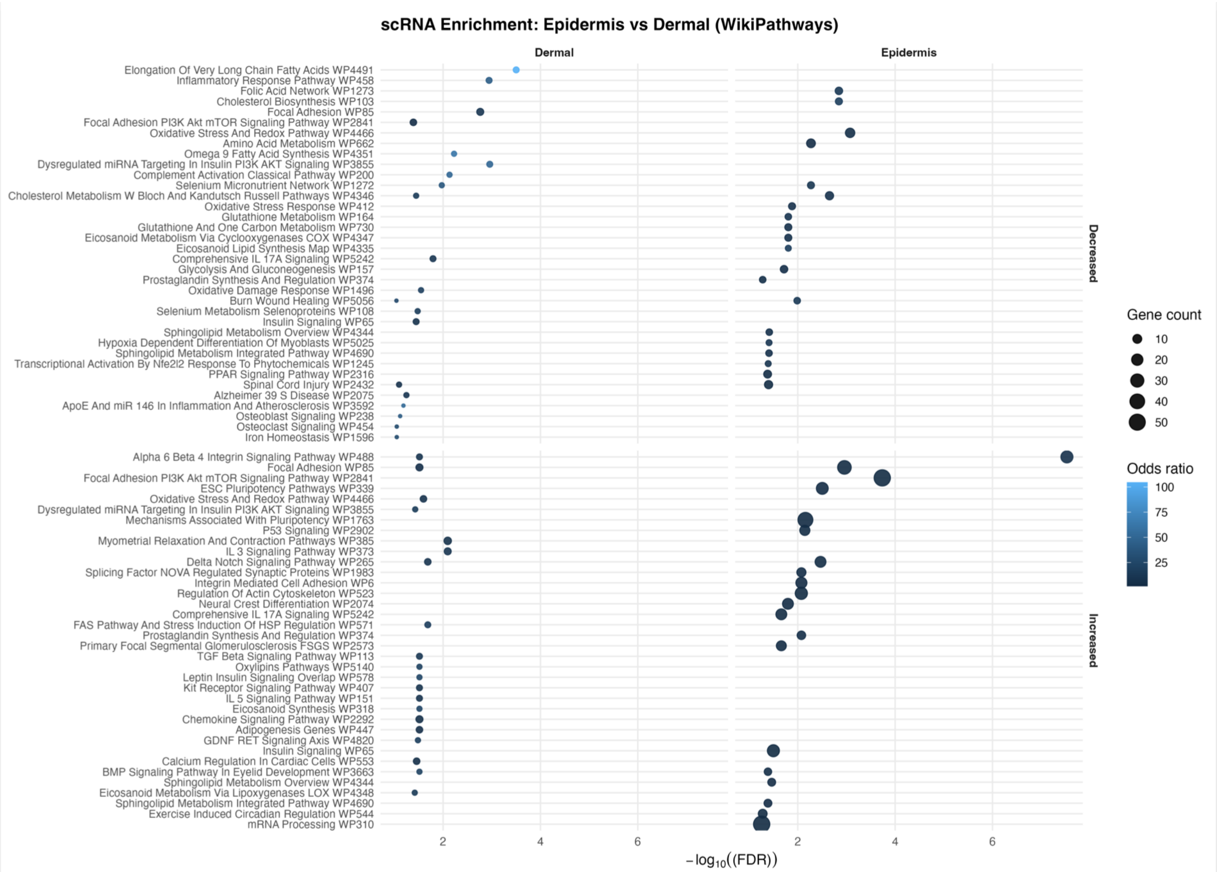


**Supplementary Figure 5. WikiPathways enrichment highlights lipid, redox, and metabolic programs engaged during disease resolution in epidermal cells.** WikiPathways enrichment analysis of differentially expressed genes identified by single-cell RNA-seq, stratified by epidermal and dermal skin compartments in KC-Tie2 mice under restored versus diseased conditions. Enriched pathways are shown separately for genes decreased (top) or increased (bottom) during disease resolution. Dot position reflects enrichment significance (-$\log_{10}$ FDR), dot size indicates the number of genes contributing to each pathway, and color denotes odds ratio. Epidermal compartments show preferential enrichment of pathways related to fatty acid and lipid metabolism, redox and oxidative stress responses, and metabolic signaling, complementing Gene Ontology and Reactome analyses shown in Supplementary Figures 3 and 4. Full WikiPathways enrichment statistics and gene lists are provided in Supplementary Tables 22-25.

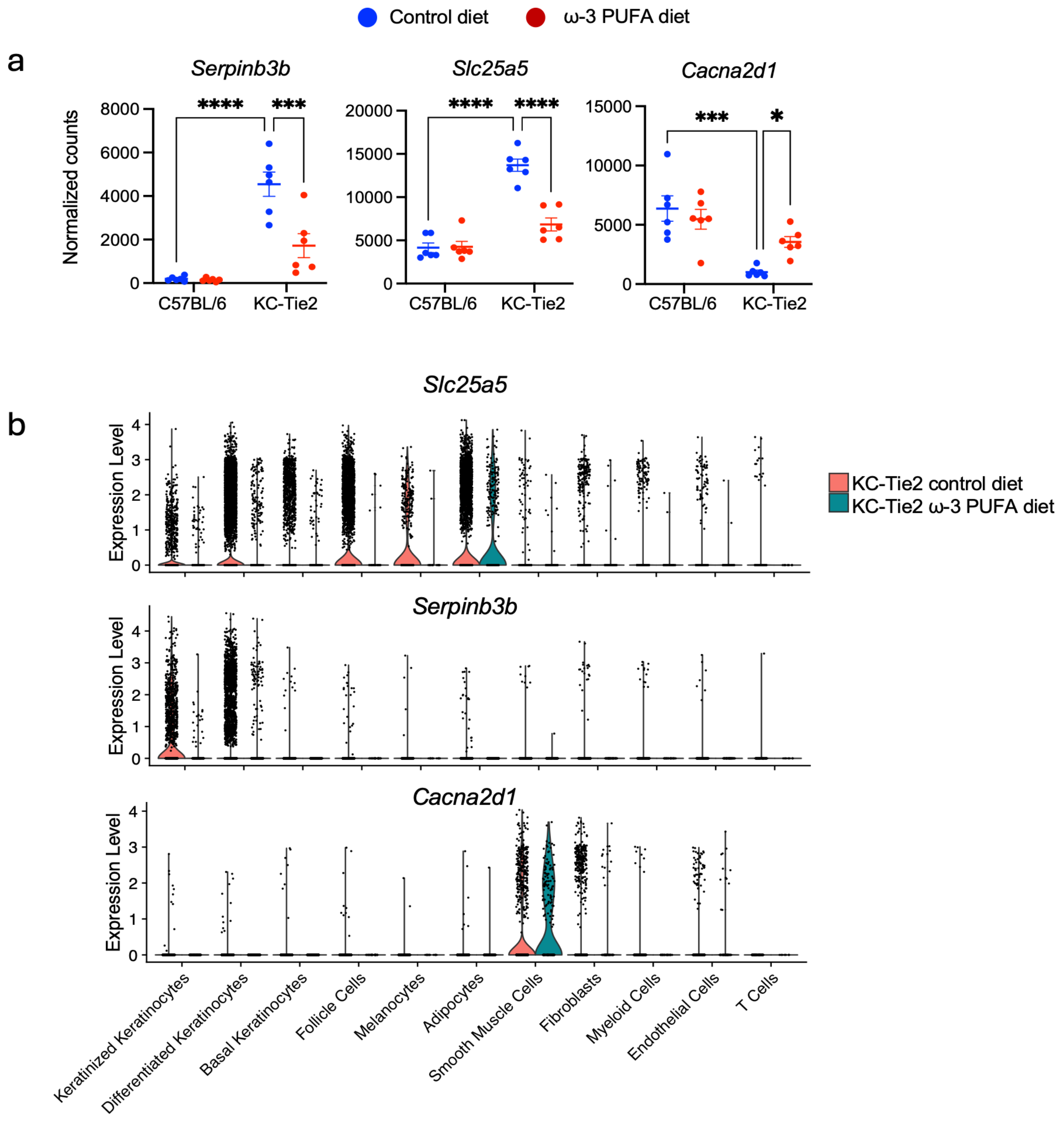


**Supplementary Figure 6. Expression of *Serpinb3b, Slc25a5*, and *Cacna2d1* across genotypes, diets, and cellular compartments. a,** Normalized bulk RNA-seq counts for *Serpinb3b, Slc25a5,* and *Cacna2d1* in dorsal skin from C57BL/6 and KC-Tie2 mice maintained on control or ω-3 PUFA diets for 6 weeks (n = 6 mice per group). Points represent individual animals; horizontal bars indicate mean ± s.e.m. Statistical comparisons were performed using two-way ANOVA with Tukey's post hoc test. *$P < 0.05$, ***$P < 0.001$, ****$P < 0.0001$. **b,** Single-cell RNA-seq violin plots showing expression of *Slc25a5, Serpinb3b*, and *Cacna2d1* across major skin cell populations from KC-Tie2 mice under control and ω-3 PUFA diets. Violin width reflects cell density; overlaid points represent individual cells. These data illustrate cell-type-specific expression patterns and diet-associated shifts without threshold filtering.

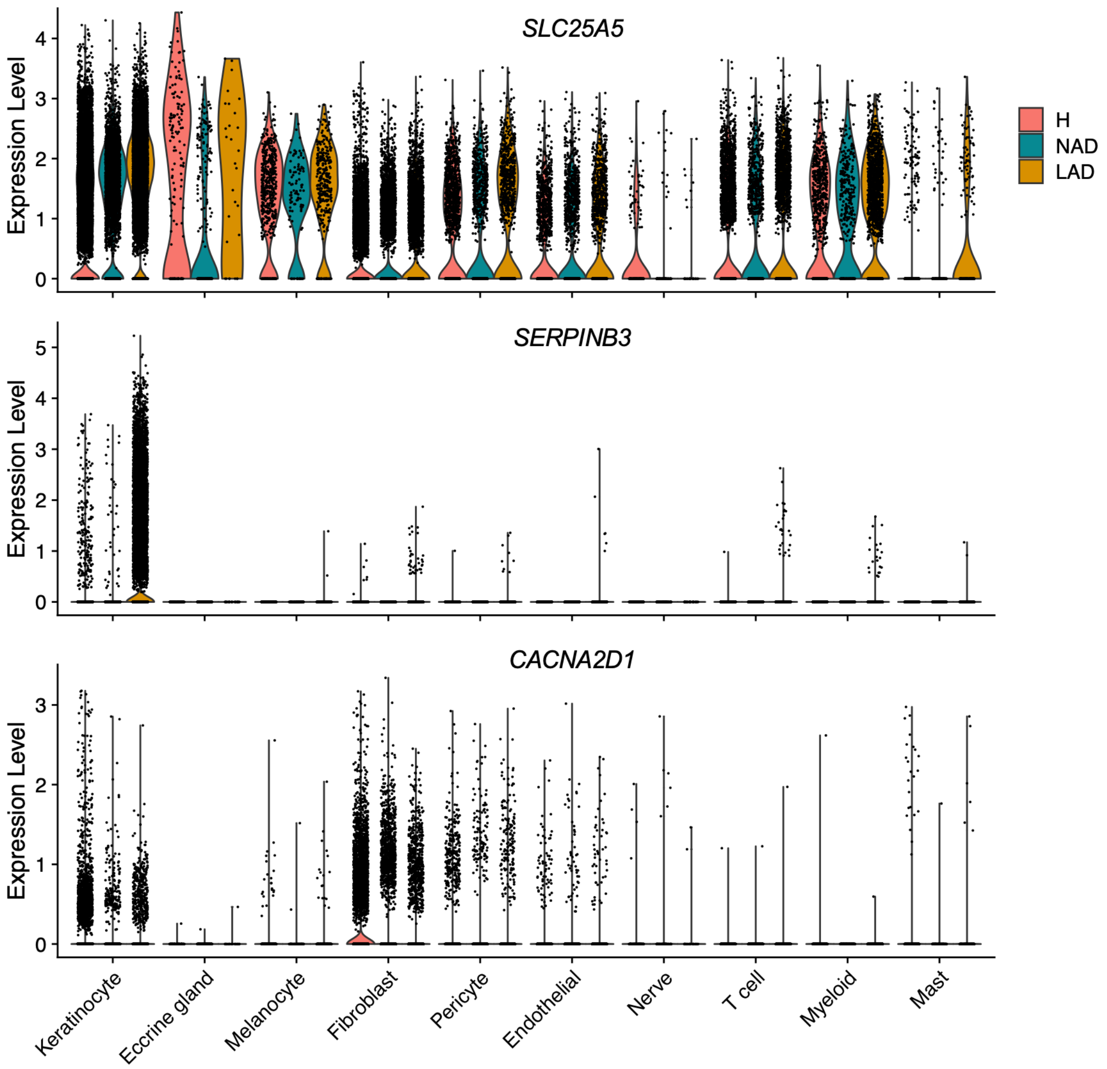


**Supplementary Figure 7. Single-cell RNA-seq expression of *SLC25A5*, *SERPINB3*, and *CACNA2D1* in healthy and atopic dermatitis human skin.** Violin plots show single-cell RNA-seq expression levels of *SLC25A5, SERPINB3*, and *CACNA2D1* across major skin cell populations from healthy volunteer skin (H), nonlesional atopic dermatitis skin (NAD), and lesional atopic dermatitis skin (LAD). Violin width reflects cell density; overlaid points represent individual cells. In lesional atopic dermatitis skin, *SLC25A5* and *SERPINB3* expression are increased in keratinocyte populations relative to healthy and nonlesional skin, whereas *CACNA2D1* expression is reduced, consistent with directionality observed in KC-Tie2 mouse skin during chronic inflammation and resolution.

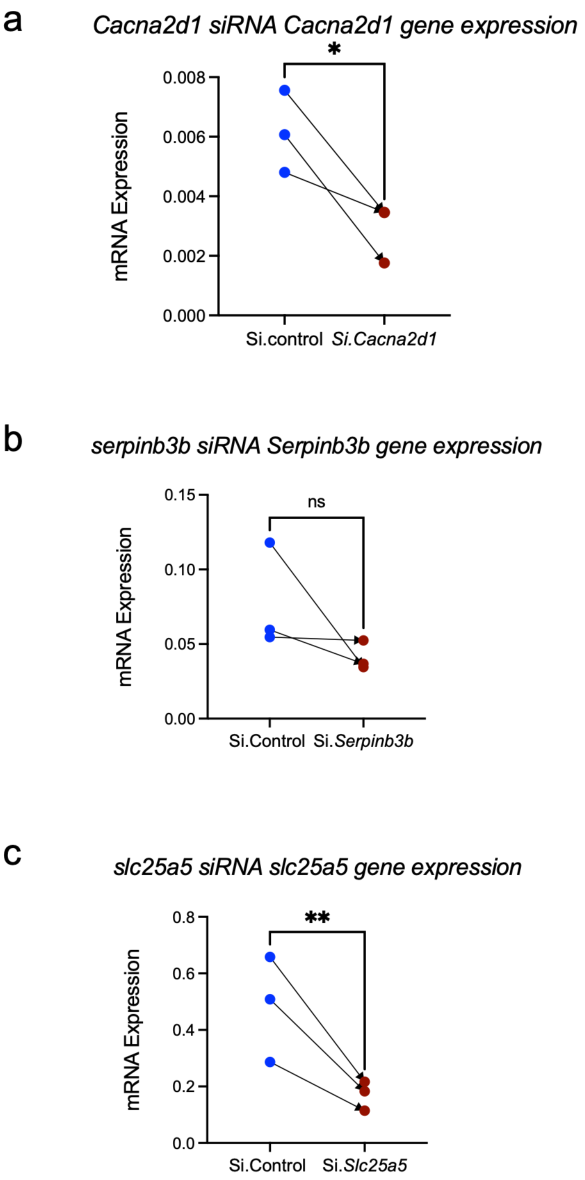


**Supplementary Figure 8. Validation of siRNA-mediated target gene knockdown in KC-Tie2 skin.** A subset of biological replicates from the KC-Tie2 siRNA perturbation experiments was randomly selected for quantitative RT–PCR analysis to confirm target gene silencing. (a-c) mRNA expression levels of **a,** *Cacna2d1*, **b,** *Serpinb3b*, and **c,** *Slc25a5* in ear skin following topical treatment with non-targeting control siRNA (si.Control) or gene-specific siRNA (si.*Cacna2d*1, si.*Serpinb3b*, or si.*Slc25a5*). Each data point represents an individual mouse; paired measurements are connected by lines. Data are shown for $n = 3$ mice per group. Statistical significance was assessed using paired Student's *t*-tests; $P < 0.05$ (*), $P < 0.01$ (**), or as indicated; ns, not significant.

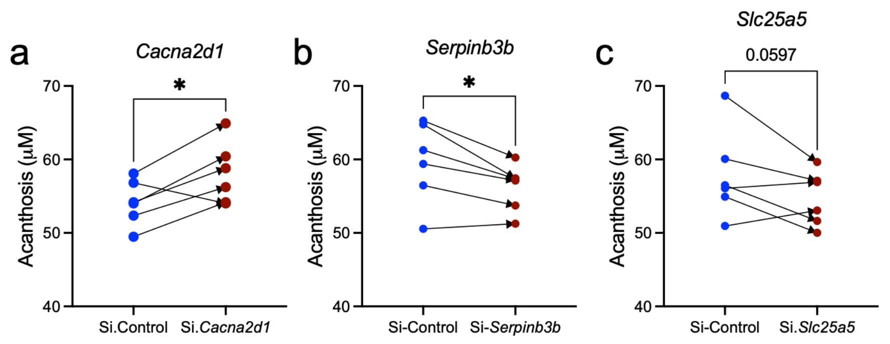


**Supplementary Figure 9. Quantification of epidermal acanthosis following siRNA-mediated gene silencing.** Epidermal thickness (acanthosis) was quantified in ear skin sections from KC-Tie2 mice following topical treatment with non-targeting control siRNA (si.Control) or gene-specific siRNA **a,** *Cacna2d1*, **b,** *Serpinb3b*, and **c,** *Slc25a5*. Acanthosis measurements (µm) were obtained from histological sections for each mouse; paired measurements from the same animal are connected by lines. Each data point represents an individual mouse ($n$ = 6 per group). Statistical significance was assessed using paired Student's $t$-tests; $P < 0.05$ (*), or exact $P$ value as indicated.

### Supplementary Table Descriptions

#### Supplementary Table 1. DESeq2-normalized bulk RNA-seq expression values

DESeq2-normalized gene expression values (ddsHTSeq_normalized) for dorsal skin samples from C57BL/6 (Bl6) and KC-Tie2 (Tie2) mice maintained on either control chow (Teklad) or ω-3 PUFA-enriched (FO) diets. Column headers encode genotype, diet, and unique biological sample identifiers. This dataset was used for principal component analysis and visualization. Differential expression testing was performed in DESeq2 using gene-level count matrices (see Methods), and genome-wide differential expression statistics are provided in Supplementary Table 2.

#### Supplementary Table 2. Genome-wide differential expression statistics for bulk RNA-seq

$Log_2$ fold change, nominal and adjusted P values (Benjamini–Hochberg FDR), and expression statistics for all detected genes across experimental contrasts. These data enable independent filtering and visualization using user-defined thresholds (as applied for downstream analyses in this study). Downstream analyses in this study used adjusted $P < 0.05$ and $|log2FC| \geq 1.5$ unless otherwise stated.

#### Supplementary Table 3. Summary of differential expression statistics across experimental comparisons

Genome-wide counts of differentially expressed genes (DEGs) across experimental contrasts, including directionality of change and non-significant genes. These summary statistics underlie the volcano plot visualizations and intersection analyses presented in Figures 2 and 3.

**Supplementary Table 4. KEGG pathway enrichment for genes increased in KC-Tie2 skin**

KEGG pathway enrichment analysis of genes significantly upregulated in KC-Tie2 skin compared with C57BL/6 controls under control diet conditions ($|\log_2$ fold change$| \geq 1.5$, FDR < 0.05). The table lists enriched KEGG pathways, nominal and adjusted P values, odds ratios, combined scores, and the contributing gene sets. These data correspond to disease-associated transcriptional programs elevated in psoriasiform skin and form the basis for pathway analyses shown in Figure 2c (upper).

**Supplementary Table 5. KEGG pathway enrichment for genes decreased in KC-Tie2 skin**

KEGG pathway enrichment analysis of genes significantly downregulated in KC-Tie2 skin compared with C57BL/6 controls under control diet conditions ($|\log_2$ fold change$| \geq 1.5$, FDR < 0.05). The table includes enriched pathways, statistical significance metrics, enrichment scores, and gene membership. These pathways represent epithelial structural, signaling, and metabolic programs suppressed in psoriasiform disease and are referenced in Figure 2c (lower).

**Supplementary Table 6. KEGG pathway enrichment for ω-3 PUFA-responsive genes decreased in KC-Tie2 skin**

KEGG pathway enrichment analysis of genes significantly decreased in KC-Tie2 skin following ω-3 PUFA exposure relative to control-fed KC-Tie2 mice ($|\log_2$ fold change$| \geq 1.5$, FDR < 0.05). Listed pathways, enrichment statistics, and gene sets correspond to disease-associated inflammatory and stress programs attenuated during resolution. These data support pathway-level analyses shown in Figure 2d (upper).

**Supplementary Table 7. KEGG pathway enrichment for ω-3 PUFA-responsive genes increased in KC-Tie2 skin**

KEGG pathway enrichment analysis of genes significantly increased in KC-Tie2 skin following ω-3 PUFA exposure relative to control-fed KC-Tie2 mice ($|\log_2$ fold change$| \geq 1.5$, FDR < 0.05). The table includes enriched KEGG pathways, statistical metrics, enrichment scores, and contributing genes. These pathways represent restored epithelial signaling, metabolic, and regulatory programs during disease normalization and correspond to Figure 2d (lower).

**Supplementary Table 8. Gene-to-axis mapping for curated metabolic-mechanical programs used in Figure 3**

Gene universe and annotation backbone for the five curated metabolic–mechanical axes in Figure 3, including supporting pathway terms and source lists and standardized mouse gene symbols used for program-level analyses.

**Supplementary Table 9. Gene Ontology (GO) Biological Process enrichment for ω-3 PUFA-responsive genes increased in KC-Tie2 skin**

Gene Ontology (GO) Biological Process enrichment analysis of genes significantly increased in KC-Tie2 skin following ω-3 PUFA exposure relative to control-fed KC-Tie2 mice (FDR < 0.05; $|\log_2$ fold change$| \geq 1.5$). The table lists enriched GO Biological Process terms, nominal and adjusted P values, odds ratios, combined scores, and contributing gene sets. These data underlie the GO enrichment analysis shown in Figure 3b and highlight coordinated regulation of cytoskeletal organization, calcium and ion handling, and metabolic and mitochondrial processes during disease resolution.

**Supplementary Table 10. Reactome pathway enrichment for ω-3 PUFA-responsive genes increased in KC-Tie2 skin**

Reactome pathway enrichment analysis of genes significantly increased in KC-Tie2 skin following ω-3 PUFA exposure relative to control-fed KC-Tie2 mice (FDR < 0.05; $|\log_2$ fold change$| \geq 1.5$). The table includes enriched Reactome pathways, statistical significance metrics, enrichment scores, and contributing gene sets. These pathways correspond to coordinated restoration of calcium-linked signaling, growth factor- and stress-responsive pathways, mitochondrial processes, and cytoskeletal and junctional organization, as visualized in Figure 3c.

**Supplementary Table 11. Single-cell RNA-seq dataset composition and cell-type distribution**

Summary of single-cell RNA-seq dataset characteristics from KC-Tie2 skin under diseased and restored conditions. The table reports total cell numbers, cell-type annotations, and relative cell-type proportions (%) for each experimental group, providing descriptive context for downstream differential expression and pathway analyses shown in Figure 4.

**Supplementary Table 12. Differential gene expression statistics across skin cell types in single-cell RNA-seq**

Comprehensive differential expression results from single-cell RNA-seq comparing restored versus diseased KC-Tie2 skin across all annotated skin cell types. For each gene and cell type, the table includes average $\log_2$ fold change, adjusted P value, and statistical significance metrics. This table provides the complete gene-level data underlying Figures 4b–d.

**Supplementary Table 13. Cell-type-specific DEG counts underlying Figure 4a**

Numeric summary of significantly differentially expressed genes (adjusted $P < 0.05$) identified by single-cell RNA-seq across major skin cell types in restored versus diseased KC-Tie2 skin. The table reports total, upregulated, and downregulated DEG counts per cell type and corresponds directly to the bar plots shown in Figure 4a.

**Supplementary Table 14. Gene Ontology Biological Process enrichment for genes increased in epidermal cells during disease resolution**

Gene Ontology (GO) Biological Process enrichment analysis of genes significantly increased in epidermal cell populations in restored versus diseased KC-Tie2 skin, derived from single-cell RNA-seq data. The table includes enriched GO terms, statistical significance measures, enrichment scores, and contributing gene sets. These results correspond to Supplementary Figure 3 (lower panel).

**Supplementary Table 15. Gene Ontology Biological Process enrichment for genes decreased in epidermal cells during disease resolution**

GO Biological Process enrichment analysis of genes significantly decreased in epidermal cell populations in restored versus diseased KC-Tie2 skin. The table reports enriched biological processes, statistical metrics, and contributing gene lists and corresponds to Supplementary Figure 3 (upper panel).

**Supplementary Table 16. Gene Ontology Biological Process enrichment for genes increased in dermal cells during disease resolution**

GO Biological Process enrichment analysis of genes significantly increased in dermal cell populations in restored versus diseased KC-Tie2 skin, derived from single-cell RNA-seq. The table includes enriched terms, enrichment statistics, and gene contributors, supporting dermal pathway analyses shown in Supplementary Figure 3.

**Supplementary Table 17. Gene Ontology Biological Process enrichment for genes decreased in dermal cells during disease resolution**

GO Biological Process enrichment analysis of genes significantly decreased in dermal cell populations in restored versus diseased KC-Tie2 skin. Enriched biological processes, statistical significance metrics, and gene lists are provided and correspond to Supplementary Figure 3.

**Supplementary Table 18. Reactome pathway enrichment for genes increased in epidermal cells during disease resolution**

Reactome pathway enrichment analysis of genes significantly increased in epidermal cell populations in restored versus diseased KC-Tie2 skin. The table includes enriched pathways, enrichment statistics, and contributing gene sets, corresponding to Supplementary Figure 4 (lower panel).

**Supplementary Table 19. Reactome pathway enrichment for genes decreased in epidermal cells during disease resolution**

Reactome pathway enrichment analysis of genes significantly decreased in epidermal cell populations during disease resolution. Enriched pathways, statistical metrics, and gene contributors are provided and correspond to Supplementary Figure 4 (upper panel).

**Supplementary Table 20. Reactome pathway enrichment for genes increased in dermal cells during disease resolution**

Reactome pathway enrichment analysis of genes significantly increased in dermal cell populations in restored versus diseased KC-Tie2 skin. The table supports dermal compartment analyses shown in Supplementary Figure 4.

**Supplementary Table 21. Reactome pathway enrichment for genes decreased in dermal cells during disease resolution**

Reactome pathway enrichment analysis of genes significantly decreased in dermal cell populations during disease resolution, with enrichment statistics and gene lists corresponding to Supplementary Figure 4.

**Supplementary Table 22. WikiPathways enrichment for genes increased in epidermal cells during disease resolution**

WikiPathways enrichment analysis of genes significantly increased in epidermal cell populations in restored versus diseased KC-Tie2 skin. The table reports enriched pathways, enrichment scores, statistical significance metrics, and contributing genes and corresponds to Supplementary Figure 5 (lower panel).

**Supplementary Table 23. WikiPathways enrichment for genes decreased in epidermal cells during disease resolution**

WikiPathways enrichment analysis of genes significantly decreased in epidermal cell populations during disease resolution. Enriched pathways and gene contributors correspond to Supplementary Figure 5 (upper panel).

**Supplementary Table 24. WikiPathways enrichment for genes increased in dermal cells during disease resolution**

WikiPathways enrichment analysis of genes significantly increased in dermal cell populations in restored versus diseased KC-Tie2 skin, supporting dermal pathway analyses shown in Supplementary Figure 5.

**Supplementary Table 25. WikiPathways enrichment for genes decreased in dermal cells during disease resolution**

WikiPathways enrichment analysis of genes significantly decreased in dermal cell populations during disease resolution. The table includes enriched pathways, statistical metrics, and gene lists corresponding to Supplementary Figure 5.

**Supplementary Table 26. Protein-coding neighbors of *Cacna2d1, Slc25a5*, and *Serpinb3b***

Protein-coding transcripts assigned to triad-centered transcriptional neighborhoods used to generate the Fig. 5D heatmap. For each gene, the table reports neighborhood assignment (*Cacna2d1, Slc25a5*, or *Serpinb3b*), the gene-triad Pearson correlation coefficient for each triad gene (*Cacna2d1, Slc25a5, Serpinb3b*), and the corresponding p-values computed across the bulk RNA-seq dataset. *Unannotated_like* flags predicted/poorly annotated loci; these were excluded from the Fig. 5D heatmap but are retained in the table for transparency.

**Supplementary Table 27. Pathway enrichment results for triad-centered transcriptional neighborhoods (GO BP, KEGG, Reactome, WikiPathways)**

Enrichment analysis results for the triad-centered neighborhoods (*Cacna2d1, Slc25a5, Serpinb3b*) using the top co-varying protein-coding neighbors for each triad gene (as defined in Supplementary Table 26). Results are reported across multiple libraries (GO Biological Process, KEGG Mouse, KEGG Human, Reactome, and WikiPathways), including term name, nominal p-value, adjusted p-value, odds ratio, combined score, and contributing genes. These raw enrichment outputs were used as input to derive the higher-order consensus themes summarized in Figure 5E and Supplementary Table 28.

**Supplementary Table 28. Consensus pathway themes derived from triad-centered neighborhood enrichment**

Higher-order functional themes compiled from the multi-library enrichment results in Supplementary Table 27 for each triad-centered neighborhood (*Cacna2d1, Slc25a5, Serpinb3b*). For each neighborhood-theme pair, the table reports: (i) libraries_supporting, the number of enrichment libraries with ≥1 enriched term mapping to the theme; (ii) max_strength, the maximum enrichment strength observed among mapped terms (as defined in the analysis pipeline); and (iii) n_terms, the total number of enriched terms assigned to the theme. These values were used to generate the consensus theme visualization in Figure 5E.

**Supplementary Table 29. $Log_2$ fold-change values underlying cytokine and inflammatory gene expression in Fig. 6c**

$Log_2$ fold-change values for selected inflammatory and keratinocyte-derived immune mediators following siRNA-mediated knockdown of *Cacna2d1, Serpinb3b*, or *Slc25a5 i*n KC-Tie2 skin. Values represent normalized qRT-PCR expression changes relative to non-targeting siRNA controls and correspond to the heatmap shown in Figure 6c. Genes shown include cytokines, chemokines, and alarmins associated with IL-23/IL-17 signaling and epidermal inflammation.

**Supplementary Table 30. Program-level correlation matrix underlying Figure 6D.**
Matrix of Pearson correlation coefficients linking expression of triad genes (*Cacna2d1, Serpinb3b, Slc25a5*) with curated epidermal differentiation, metabolic, inflammatory, and structural gene programs derived from bulk RNA-seq data. Values shown correspond to the data visualized in Figure 6D.
